# Structure, motion, and multiscale search of traveling networks

**DOI:** 10.1101/2024.01.16.575883

**Authors:** Nate J. Cira, Morgan L. Paull, Shayandev Sinha, Fabio Zanini, Eric Yue Ma, Ingmar H. Riedel-Kruse

## Abstract

Network models are widely applied to describe connectivity and flow in diverse systems. In contrast, the fact that many connected systems move through space as the result of dynamic restructuring has received little attention. Therefore, we introduce the concept of ‘traveling networks’, and we analyze a tree-based model where the leaves are stochastically manipulated to grow, branch, and retract. We derive how these restructuring rates determine key attributes of network structure and motion, enabling a compact understanding of higher-level network behaviors such as multiscale search. These networks self-organize to the critical point between exponential growth and decay, allowing them to detect and respond to environmental signals with high sensitivity. Finally, we demonstrate how the traveling network concept applies to real-world systems, such as slime molds, the actin cytoskeleton, and human organizations, exemplifying how restructuring rules and rates in general can select for versatile search strategies in real or abstract spaces.

## MAIN

Network models play a key role in representing and understanding the characteristics of complex connected systems. Networks consist of nodes connected by edges, and networks can be generated in different ways, for example, by randomly [1], regularly, or semi-randomly [2–4] connecting existing nodes to each other, or by iteratively branching from an initial seed [5–12]. Processes of interest can often be understood using only the connectivity information between nodes [13]; however, additional constraints arise in spatial networks where each node is also embedded inside a real or abstract space [14]. Networks can be static or dynamic [15], and movement in the context of networks is usually discussed in terms of processes occurring on the network, for example, an agent walking over a network [16, 17] or information or disease spreading through a network [18, 19], or in terms of a network developing from a rooted position like neuronal dendrites [12] or branched organs [20]. Despite significant work on temporal networks [15], surprisingly, there appears to be little systematic work on how networks themselves travel through space away from an initial location *without* a permanent anchor point.

### ‘Traveling networks’

We therefore introduce the concept of ‘traveling networks’, i.e., connected systems that change their location in space over time by rearranging their structure. This class of networks is motivated by various real-world systems, for example: (1) The slime mold *Physarum polycephalum* traverses the environment in search for food by rearranging its branched network (figure 1a, supplementary movie 1) [21, 22]. (2) The subcellular actin cytoskeleton network drives eukaryotic cell movement through polymerization and depolymerization of its filaments (figure 1b) [23–25]. (3) Human corporations move through high-dimensional market spaces by creating new products and phasing out older ones (figure 1c) [26–30]. For each of these systems, a characteristic time scale *τ*_*r*_ exists after which none of their original structure remains and the network occupies a distinct new region of space (figure 1a-c). This makes these networks distinct from other well-studied networks that are able to dynamically grow and branch but do not leave their original rooted location [5, 7–12].

**Figure 1:**
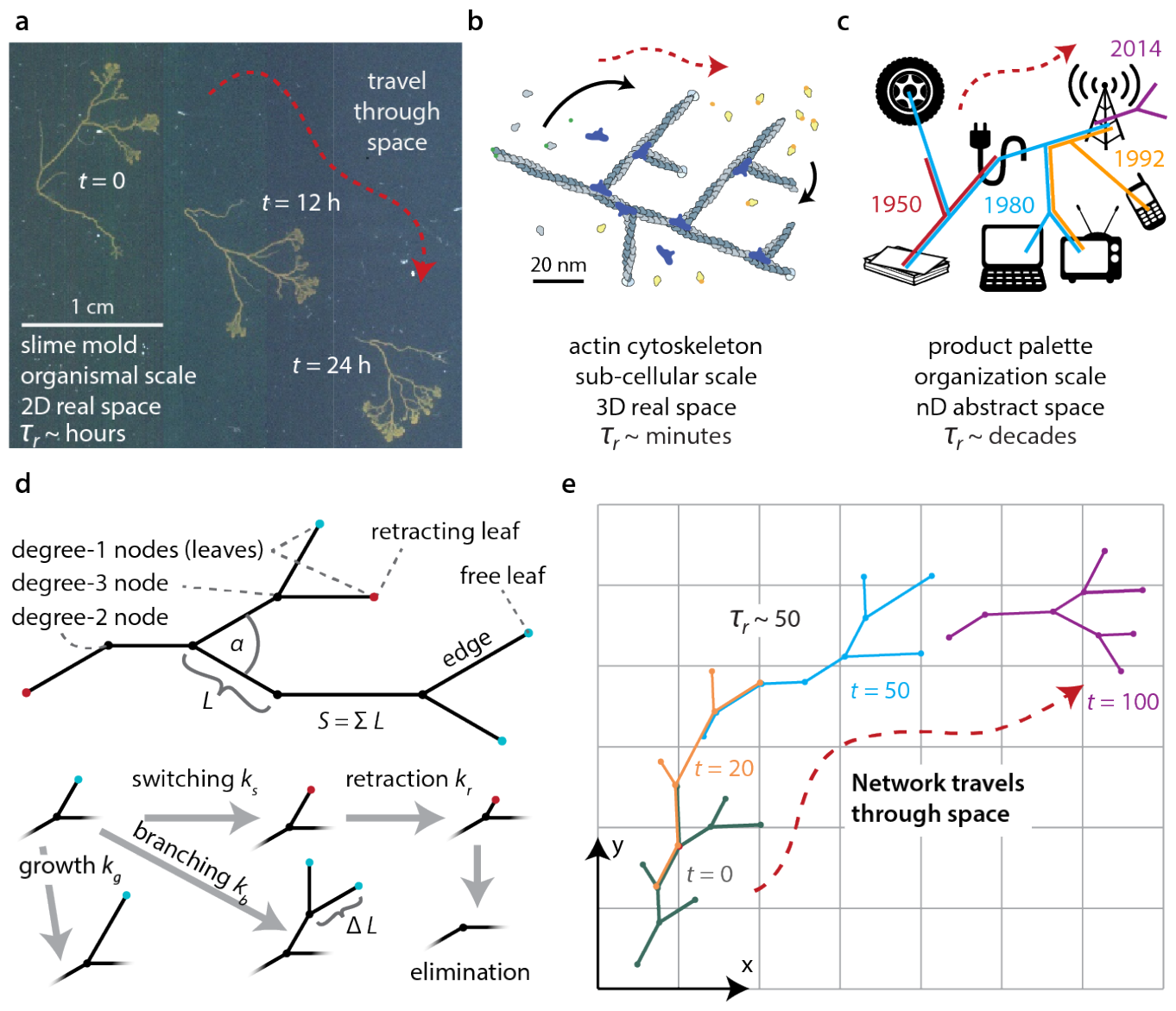
‘Traveling networks’ that actively traverse space via dynamic restructuring are found in diverse contexts. **a-c)** Examples of traveling networks: **a)** slime molds (2D real space) [21], **b)** actin cytoskelton (3D real space) [23], **c)** human organizations (high dimensional abstract space, depicting select products of Nokia Corporation over time) [31–33]. **d)** Model of a traveling network based on an acyclic binary tree with two leaf types. ‘Free leaves’ (cyan) branch at rate *k*_*b*_, grow at rate *k*_*g*_, and switch at rate *k*_*s*_, to become ‘retracting leaves’ (red) which retract at rate *k*_*r*_ until they reach a degree-three node. The size of the network *S* is the sum of the edge lengths *L*. **e)** The model from (d) results in a traveling network with conceptual similarity to (a-c). Time in arbitrary units. (a-c, e: Red arrow illustrates how networks travel through space; *τ*_*r*_ represents typical time scale after which the network has completely remodeled itself.) (Image b adapted from [23, 34].)

### A traveling network model

This motivates a deeper investigation into the constraints and opportunities experienced by such traveling networks. Here we propose and analyze a tractable model (figure 1d) that involves a tree with maximum degree (number of edges connected to a node) of three. The leaves (degree-one nodes) are stochastically manipulated and can be in either a ‘free’ or a ‘retracting’ state. Free leaves undergo three possible manipulations: 1) growth by adding unit length Δ*L* at rate *k*_*g*_; 2) branching at rate *k*_*b*_, by creating two new free leaves - each with an edge of length Δ*L* at an angle ±*α/*2 from the previous edge, thereby converting the original leaf into a degree-3 node; and 3) switching by becoming a retracting leaf at rate *k*_*s*_. Retracting leaves undergo only one manipulation: removing length Δ*L* at rate *k*_*r*_ until they reach a degree-three node, upon which they are eliminated and generate a degree-two node. In this model, retracting leaves cannot switch back to being a free leaf. The amount of resources available to the system is represented by the overall size of the network, *S*, which is defined as the sum of all the edge lengths and implies that more spatially distant nodes require more resources to connect. These rules lead to networks that travel through space due to restructuring as illustrated in figure 1e (supplementary movies 2 and section 3). This model clearly neglects many aspects of any particular real system (figure 1a-c), yet it proves to be very informative for analyzing many key properties and behaviors of general importance to traveling networks, as demonstrated in the following.

### Network structure and criticality

Under steady state conditions, these networks operate at the critical point between exponential increase and decrease of free leaves, *k*_*b*_ = *k*_*s*_, resembling a critical Galton-Watson process [35]. Nondimensionalizing the time and length scales makes it possible to express the steady state model with only two truly independent parameters (apart from the branching angle *α*): the total number of edges *N*_*E*_ and the rescaled branching rate 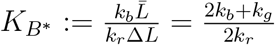 All networks sharing these parameters then belong to the same ‘universal form’ and show equivalent rescaled behavior (supplementary section 4.5.2 and 4.6). Since many real-world systems likely have more direct control over their specific rates than over *N*_*E*_ and *K*_*B\**_, we instead focus the following analysis on a representation that is based on network size *S* and the original rates (*k*_*b*_, *k*_*g*_, *k*_*r*_) or rates non-dimensionalized by *k*_*r*_ (*K*_*B*_ := *k*_*b*_*/k*_*r*_, *K*_*G*_ := *k*_*g*_*/k*_*r*_).

We implemented a stochastic simulation embedded in 2D space to investigate and visualize the model’s behavior, though much of the following analysis generalizes to higher dimensions. We set *K*_*B*_ = *K*_*S*_ and numerically impose conservation of size *S* (supplementary section 8.1). Arbitrary starting configurations converge to steady state dynamics where branching, growing, and retracting leaves cause the network to travel through space (supplementary movie 2). The network structure varies characteristically with *K*_*B*_ and *K*_*G*_ as indicated by the number of retracting (*N*_*R*_), free (*N*_*F*_), and total (*N*_1_ = *N*_*F*_ + *N*_*R*_) leaves (figure 2a-d, supplementary movie 3). Networks with high *K*_*B*_ have more retracting leaves; if both *K*_*G*_ and *K*_*B*_ are small, most leaves are free, and a large ratio of *K*_*G*_ : *K*_*B*_ results in longer edges, fewer nodes, and fewer total leaves.

**Figure 2:**
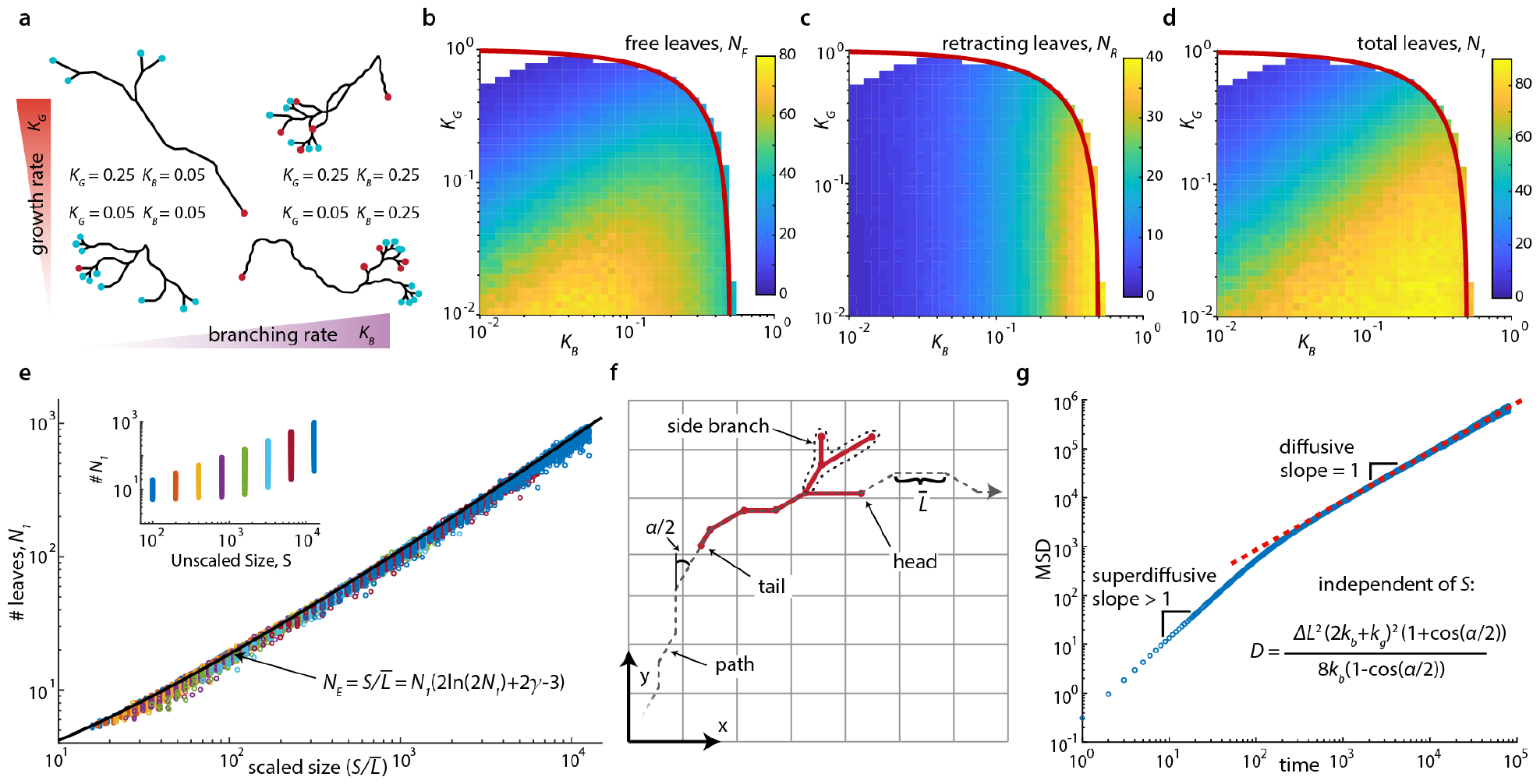
Network structure and movement are determined by restructuring rates. **a)** A range of network structures are possible, illustrated by four typical networks at high/low *K*_*B*_/*K*_*G*_ having different edge lengths, numbers of free leaves (cyan) and retracting leaves (red), and leaf type ratios (simulation of model from figure 1d; size, *S* = 70, supplementary movie 3). **b-d)** *K*_*B*_ and *K*_*G*_ regulate the structure, exhibited by the mean numbers of free leaves (b), retracting leaves (c), and the total leaves (d) (simulated networks of *S* = 800; red lines: 2*K*_*B*_ + *K*_*G*_ ≤ 1 boundary of possible steady state behavior). **e)** Total leaves, *N*_1_, for various sizes, *S*, across the steady-state parameter space of *K*_*B*_ and *K*_*G*_ shown in (b-d). Number of leaves depends primarily on the average branch length 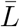 as captured by the relation *S* = *N*_1_(2 ln(2*N*_1_) + 2*γ* − 3), where *γ* is the Euler-Mascheroni constant (black line). Inset: *N*_1_ does not collapse with unscaled *S*. **f)** The longer term network movement reveals a ‘tail’ and, in retrospect, a unique ‘path’, and a “head” (supplementary movie 4). **g)** The network (center of mass) moves superdiffusively over short time scales and diffusively over long time scales, with the diffusivity set by *k*_*b*_, *k*_*g*_, Δ*L*, and *α. k*_*b*_ = 0.2, *k*_*g*_ = 0.1, *α* = 60, *S* = 200 shown here; red dashed line indicates analytical result.

The structure of a network is characterized by the numbers of different node types and their connectivity and can be crucial to network performance. We therefore derived analytical relationships for the key steady state characteristics of these quantities in the model (supplementary section 4). In agreement with numerical results, arguments in the steady state reveal that the edge lengths are distributed geometrically with mean length, 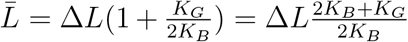. The number of all edges is *N*_*E*_ = *N*_1_ + *N*_2_ + *N*_3_ − 1 (where *N*_*i*_ is the number of nodes of degree *i*), with *N*_3_ = *N*_1_ − 2 and *N*_2_ = *N*_*E*_ + 3 − 2*N*_1_, and on average 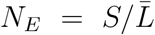). The ratio of the number of retracting to free leaves is 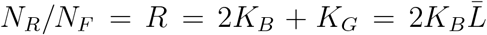. By analyzing a recursive limit case, we found the final determining formula *S* ≈ *N*_1_(2 ln(2*N*_1_) + 2*γ* − 3) (*γ* is the Euler-Mascheroni constant), which is in agreement with simulations over a wide parameter space (figure 2e, supplementary section 4.4). The branching angle *α* (figure 1d) influences the appearance of the graph when embedded in space but not the number of each node type or edge length distribution. Hence 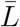, *R, N*_1_, *N*_2_, *N*_3_, *N*_*F*_, *N*_*R*_, and *N*_*E*_ are fully determined by just the three parameters *K*_*G*_, *K*_*B*_, and *S*, with 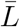 and *R* even being independent of the network size *S* (supplementary section 4).

### Network restructuring and traveling

Next, we analyzed how the network actually travels through space due to its branching, growing, retracting, and eventually disappearing leaves. One fundamental quantity for traveling networks is their relocation time, *τ*_*r*_, the time it takes for the entire network to reorganize such that it occupies a new spatial position and none of its initial edges exist anymore (figure 1a-c,e). Interestingly, we find that the relocation time depends only on the number of leaves and branching rate 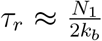 (supplementary section 5.4). Another fundamental aspect is a description of the network’s longer-term movement. In this acyclic model, for times larger than *τ*_*r*_, we find that a single unique path exists that connects the network’s initial and final positions (gray dashed line in figure 2f, supplementary movie 4). We define the ‘tail’ node as the oldest node in the network. The tail follows this unique path on the network’s trailing end, advancing at rate *k*_*r*_ and intermittently pausing whenever it reaches a degree-three node. We also define a ‘head’ (which can only be identified in retrospect) as the free leaf that moved along this path at the leading edge of the network never switching or retracting. The head advances with speed *v*_*h*_ = Δ*L*(2*k*_*b*_ + *k*_*g*_). To maintain a steady state length between head and tail along the path, 2*k*_*b*_ + *k*_*g*_ = *f* · *k*_*r*_ must hold, where *f* ≤ 1 is the fraction of time the tail is retracting rather than paused. In the framework nondimensionalized by *k*_*r*_, this implies 2*K*_*B*_ + *K*_*G*_ ≤ 1, which constrains allowable values of *K*_*B*_ and *K*_*G*_ for steady state dynamics (red lines in figure 2b-d). Hence this type of network behaves like a chain of nodes moving along a random path while extending and retracting temporary side branches.

Focusing on the head dynamics, we identified a correlated random walk, which, over longer time scales, also corresponds to the motion of the network’s center of mass. This random walk has step size 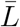, step rate 2*k*_*b*_, and turning angle *α/*2. The diffusivity of the head and correspondingly the entire network can be written entirely in terms of the fundamental parameters as 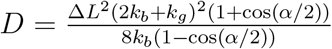 (figure 2g), which corresponds to the freely rotating chain model in polymer physics [36, 37]. Interestingly, *D* is independent of network size. Networks with high *k*_*g*_ and low *k*_*b*_ diffuse fast (large *D*) but have structures with few free leaves (figure 2b). Here a small number of free leaves might be considered a risky configuration, since stochastic elimination of all free leaves at once could lead to catastrophic collapse of the entire network (supplementary section 8). The above expressions provide quantitative insights into the network’s structure and motion, moreover, since the same rates simultaneously impact *both* structure *and* motion, this analysis highlights how traveling networks face inherent tradeoffs.

### Passive spatial search

Such tradeoffs become particularly important for spatial search, a task that many traveling networks perform in real or abstract spaces (figure 1a-c). We therefore analyzed the efficacy of random search for different target sizes by counting all unique boxes of different sizes [38] that a network occupied within a given time period (figure 3a). We ran corresponding simulations on four networks with different combinations of high and low *K*_*B*_ and *K*_*G*_ (figure 3b). The low *K*_*B*_, high *K*_*G*_ network collected the most large boxes, which can be understood through the expression for *D*, which indicates larger diffusivity for large *K*_*G*_. Networks with higher *K*_*B*_ collected the most small boxes since new lengths of 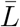 are added at rate *N*_*F*_ ·2*k*_*b*_ (before the network begins crossing itself). The low *K*_*B*_, low *K*_*G*_ network diffuses slowly and does not add much length per time hence it is not an effective searcher at any length scale. Data from all four networks actually collapses to a single curve when rescaled, corresponding to the universal form discussed earlier (figure 2b inset, supplementary sections 4.6 and 6.1). The absolute value of the slope of these plots is the Hausdorff dimension of the network history which increases toward 2 with long runtimes as self-crossing becomes more frequent (supplementary section 6.1.4), consistent with Brownian trails in two or more dimensions [38]. Hence traveling networks can tune their restructuring rates for optimal multiscale search depending on the relevant time and length scales.

**Figure 3:**
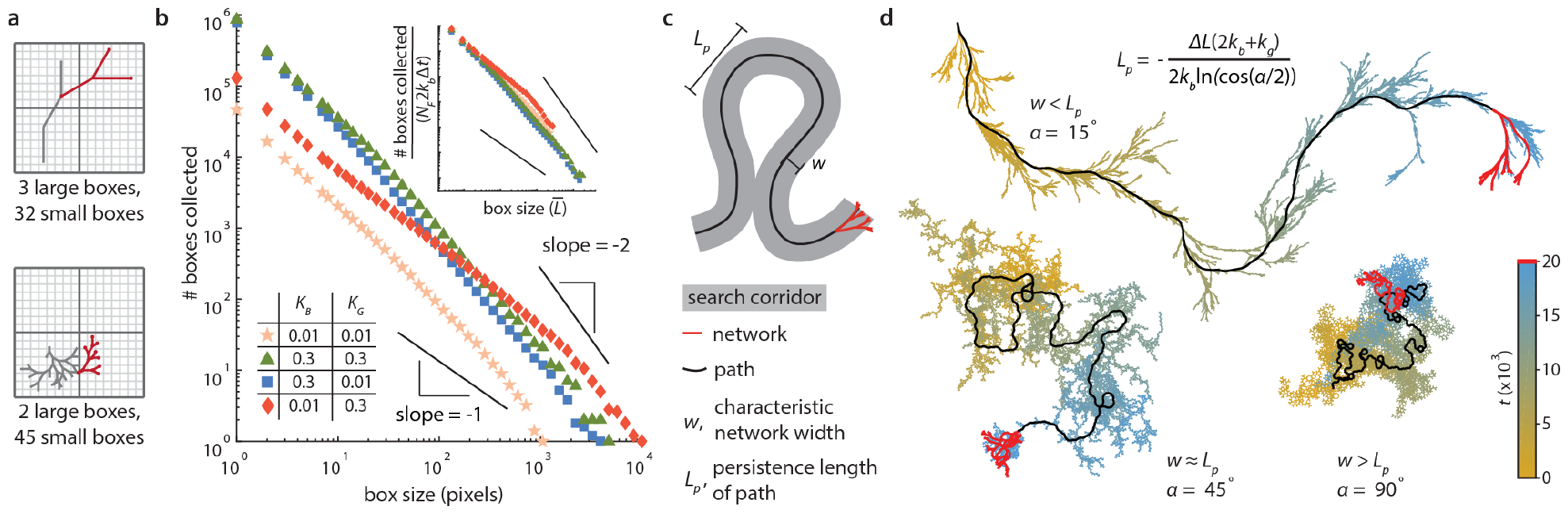
Traveling networks can control their passive search behavior across multiple length and time scales through regulation of their restructuring rates. **a)** Schematic of box counting quantification over time illustrated for two different networks with different search efficacy at different spatial resolutions (gray is the full past history, and red is final network position). **b)** Four networks, *S* = 100, runtime: Δ*t* = 2000, *K*_*B*_=(0.01, 0.3), *K*_*G*_=(0.01, 0.3) show that having low *k*_*b*_ and high *k*_*g*_ maximizes diffusivity and collection of large boxes (red diamonds), and high *k*_*b*_ optimizes for collecting small boxes (blue squares and green triangles), illustrating a tradeoff between coarse and fine search. Low *k*_*b*_ and low *k*_*g*_ (peach stars) have worse performance across all sizes. (The slopes of -1 and -2 are inserted for visual guidance.) Inset: Rescaling box size by 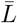 and boxes collected by *N*_*f*_ · 2*k*_*b*_ · Δt (the total expected 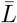 units added) collapses this data onto a single curve. **c)** Characteristic length scales of traveling networks include the network width, *w*, and path persistence length, *L*_*p*_ (supplementary sections 5.2.1 and 6.1.1). **d)** Example networks of different *w* and *L*_*p*_ generated by varying branching angle, *α* (15, 45, 90 degrees) at constant *S* = 200, *k*_*b*_ = 0.01, *k*_*g*_ = 0, *k*_*r*_ = 1, Δ*t* = 20, 000. The most recently occupied historic network positions fade from blue (recent) to gold (old); red is the final position, and black indicates the path. For *w < L*_*p*_, the network sweeps a search corridor along its path, and for *w > L*_*p*_ the network behaves as a jittering blob.

Network size and branching angle affect how much space the side branches can explore, thus we analyzed how these parameters impact search behavior. Using the above expressions and understanding of the network’s structure and motion, we find that the path has a characteristic persistence length, 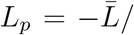 ln(cos(*α/*2)), and the network has a characteristic search width, *w*, (figure 3c) which itself has a more complex dependence on 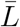, *α*, and *S* (supplementary section 6.1.1). For *w << L*_*p*_ the network behaves like a particle sweeping a corridor. For *w >> L*_*p*_ the network behaves like a jittering blob conducting a fine-scale search of limited scope. This ‘search morphology’ is sensitive to *α* as illustrated in figure 3d. Counting boxes of size 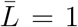 over time reveals that the optimal angle for search depends on the search duration, with *w* ≈ *L*_*p*_ collecting more boxes over shorter times and *w < L*_*p*_ collecting more boxes over longer times in these examples (supplementary section 6.1.5).

### Active and biased spatial search

Beyond the random restructuring and motion considered so far, we now consider traveling networks that respond to their environments through biased restructuring. We extended the model by making the switching rate of every free leaf increase or decrease depending on whether the environment, encoded by a scalar field, at the spatial position of each leaf is smaller or larger, respectively, compared to the field averaged across all free leaves. This leads to a biased motion of the network up a field gradient (figure 4a, supplementary movie 5, supplementary section 8.4.1). To capture environmental resource consumption as caused by natural systems (figure 1a) [21, 22], we coupled the network position to modifications of the field itself, leading to the network’s biased motion through a changing environment (figure 4b, supplementary movie 6). Many other model alterations can capture various environmental responses, for example, modifying rates based on only local information (supplementary movie 7, supplementary section 8.4.2). Importantly, these traveling networks can function as highly sensitive detectors as network size conservation self-organizes them to operate at a critical point [39, 40] where small environmentally-induced changes to their fundamental underlying restructuring rates can cause enormous changes to their emergent structure and motion through local exponential growth and decay (supplementary section 6.2).

**Figure 4:**
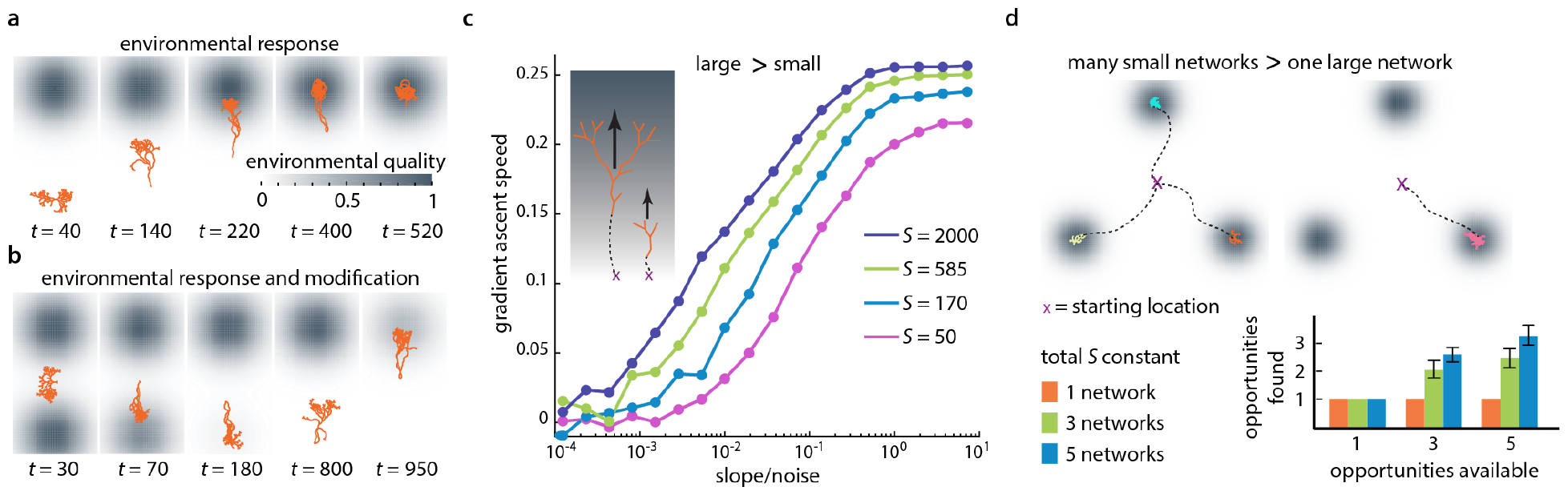
Traveling networks can control their active search behavior across multiple length and time scales through regulation of their restructuring rates. **a**,**b)** Traveling networks can perform biased search by making leaf switching rates depend on the local vs. globally sensed environment in a stationary gradient (a) or even while modifying their environment (b). **c)** When ascending noisy gradients, large networks tolerate lower slope-to-noise-ratios and ascend faster than small networks. Different network sizes: *S, α* = 60 degrees, *k*_*b*_=0.1, *k*_*g*_=0. **d)** Multiple small networks exploit more opportunities in environments with multiple opportunities than a single large network of the same total size. (n=40 trials, error bars are one standard deviation).

Many real-world traveling networks (figure 1a-c) can split into multiple smaller entities or merge into larger structures (figure 1a,c) [21, 41], suggesting that trade-offs exist between network size and network number for performing efficient search tasks. We tested this hypothesis with simulations (figure 4c,d). We found that for responsive networks moving up a linear, noisy gradient, larger networks are able to move faster and tolerate higher noise levels than smaller networks (figure 4c). However, in a landscape with different numbers of ‘opportunities’, represented by local Gaussian maxima, multiple small networks are able to capture more opportunities than a single large network of the same total size (figure 4d, supplementary movie 8). Hence it can be more advantageous to allocate the resources into one large network or into multiple smaller networks - depending on the specific environment and search task.

### Application to real-world systems

Connecting the traveling network concept and our analysis results back to our motivating examples (figure 1a-c), we note that evidence for similar connections between underlying parameters choices, network structure, and strategic movements exists in the literature (figure 5a-c). (1) Takamatsu *et al*. [8] experimentally observed that *Physarum* adopts different network structures depending on its environment. More highly branched sheet-like networks formed under favorable nutrient-rich conditions contrasting with longer, spindly networks that formed in adverse chemical environments (figure 5a) (supplementary movies 7 and 9). (2) The actin literature connects (de-) polymerization and branching rates to cytoskelatal structure and movement, which then affect the number of filopodia and higher-level (search) behaviors of cells (figure 5b) [24, 42]. There is also experimental evidence for self-organized criticality in actin networks [43] similar to our model predictions (supplementary text 4.4). (3) The business strategy literature identified a positive correlation between shareholder returns and capital reallocation among a company’s business units (figure 5c) [44, 45], suggesting that active movement through the space of market opportunities via dynamic changes (termed ‘seeding’, ‘nurturing’ and ‘pruning’) is important for productivity. Additionally, there is also analysis of which factors make it advantageous for companies to merge or split (compare to figure 4c,d) [46, 47]. In the context of a traveling network model, we identify how rates (*k*_*b*_, *k*_*g*_, *k*_*r*_) should be changed in order to select the desired behavioral strategy (figure 5a-c). Conclusions from the model analyzed here can be summarized in a concise 2D parameter space (figure 5d) (supplementary sections 4.5.2 and 4.6) that reveals various trade-offs, such as between coarse fast search vs. fine slow search or when switching between ‘exploration and exploitation’ [48] or when avoiding the risk of catastrophic collapse by operating sufficiently far away from non-steady state conditions (red line, figure 5d).

**Figure 5:**
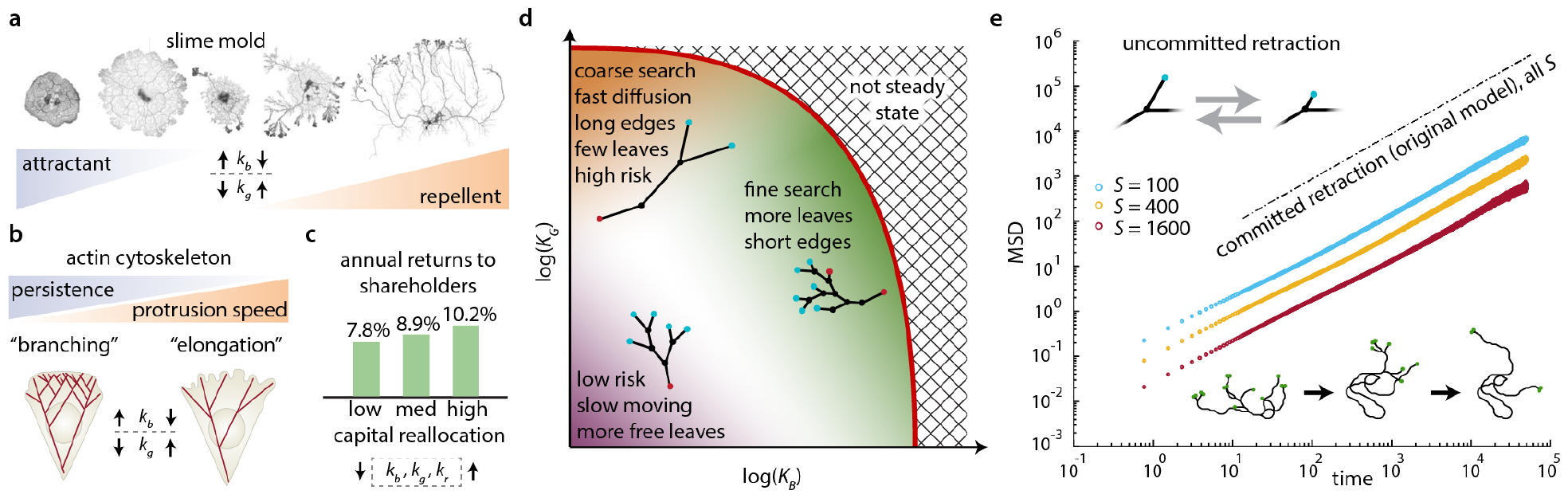
Real-world traveling networks can optimize for different behaviors through dynamic changes to their restructuring rates. **a)** Different *Physarum* network morphologies are seen in adverse and favorable environments (right vs. left) [8]. **b)** Changes in actin cytoskeleton structure and resulting cell movement due to network restructuring by branching and elongation [42]. **c)** Positive association between more capital reallocation (higher restructuring rates) and returns for corporations, 1990-2005, based on three quantiles of n=1616 companies [44]. (Images in (a-c) adapted from the corresponding references. Up and down arrows indicate how restructuring rates might be changed within a traveling network model to achieve the behaviors; rate changes deduced qualitatively from the literature [8, 42, 44, 45]. **d)** Summary of behavioral and ecologically relevant network properties and their codependence on the underlying restructuring rates in a concise 2D parameter space (example network morphologies superimposed, overall conclusions independent of *S*; colors of nodes as in figure 1d, time nondimensionalized by *k*_*r*_). **e)** A traveling network model without persistent retraction (‘uncommitted retraction’ model) explores the space more slowly than the original model (figure 1d), and becomes even less efficient with increasing network size.

### Alternative models

So far, our analysis focused on one model with particular assumptions (figure 1d), but many real-world traveling networks likely follow very different restructuring rules. As a contrasting example, we analyzed a similar model, but where retracting leaves are now also allowed to revert back into a growing and branching leaves. Hence only one leaf type exists, and leaves frequently alternate back and forth between retraction and growth (figure 5e). We found that for such an ‘uncommitted retraction’ model the resulting networks do not display the range of morphologies seen in figure 2a; instead these networks degenerate into a single chain of nearly unbranched degree-two nodes (figure 5e inset, supplementary section 7.1, supplementary movie 10). Furthermore, the diffusion rate significantly slows with increasing network size *S*, unlike the much faster and size-independent diffusion of the original model (figure 5e, figure 2g). Hence the ‘stay committed to retraction once started’ rule appears important here for achieving structural complexity as well as for effective motion and search, and, more broadly, rules as well as rates can have a profound impact on traveling network performance.

## Discussion

Here we introduced the concept of traveling networks and revealed how both restructuring rates and restructuring rules allow such systems to achieve different behaviors and strategies while traversing space. Alternative rule choices should be investigated in the future (supplementary section 7.2), promising a rich set of theoretical results and applied insights into specific systems. We believe that this traveling network concept will be helpful in understanding and optimizing various other dynamic systems, including interbreeding populations traveling through the space of genotypes [49], swarm robots restructuring their communication tree [50], or researchers and labs pursuing different strategies when traveling through ‘knowledge space’ with presumed correlations for scientific impact [48, 51].

