## Supplementary Information for "Structure, motion, and multiscale search of traveling networks"

### Supplementary Material

**1 Supplementary Material Overview** **4**

**2 Key symbols used** **5**

**3 Basic assumptions and definitions** **6**

**4 Network structure** **7**

**9 Cultivation and analysis of *Physarum*** **54**

**10 Supplementary movie descriptions** **55**

**References** **56**

### 1 Supplementary Material Overview

The supplementary materials are organized as follows: First we define the model. Then we derive various aspects of average network structure in terms of the restructuring rates and network size, using steady state, nondimensionalization, and limit case arguments. We then derive various aspects of the network's motion including diffusivity and relocation time. We then use the derived aspects of structure and motion to understand search in unbiased and biased contexts. Finally we mention alternative traveling network models and cover details of numerical implementations and methods.

### 2 Key symbols used

| symbol | description |
|---|---|
| $S$ | network size |
| $k_g$ | growth rate |
| $k_b$ | branching rate |
| $k_s$ | switching rate |
| $k_r$ | retraction rate |
| $k_d$ | rate of retracting leaf elimination |
| $k_x$ | rate of retracting leaves reaching degree-two nodes |
| $\alpha$ | branching angle |
| $\Delta L$ | change in edge length from each rearrangement |
| $N_F$ | number of free leaves |
| $N_R$ | number of retracting leaves |
| $R$ | ratio of retracting to free leaves: $\frac{N_R}{N_F}$ |
| $N_1$ | number of degree-one nodes |
| $N_2$ | number of degree-two nodes |
| $N_3$ | number of degree-three nodes |
| $N_E$ | number of edges |
| $\bar{L}$ | average edge length |
| $\bar{L}_R$ | average retraction length |
| $K_G$, $K_B$, $K_S$, $K_R$, $K_D$, $K_X$ | rates above normalized by retraction time |
| $K_{B^*}$ | branching rate in model normalized by length and time |
| $f$ | fraction of time the tail is moving |
| $\Delta t$ | runtime |
| $D$ | diffusivity of the network |
| $L_p$ | persistence length |
| $\tau_r$ | relocation time |
| $L_{SB}$ | side branch length |
| $a_{max}$ | normalized age of the oldest node |
| $w$ | characteristic network width |

### 3 Basic assumptions and definitions

The model involves a binary (acyclic) tree with nodes of degree -one, -two and -three. The leaves (degree-one nodes) are stochastically manipulated. There are two different types of leaves, free and retracting, the numbers of which are given by $N_F$ and $N_R$, respectively. Free leaves can undergo three possible manipulations: first they can grow, which involves advancing through space away from the node they are connected to; second they can branch, creating two new free leaves each at angle $\pm\alpha/2$ from the previous edge and turning the previous leaf into a degree-three node; and third they can switch states to become a retracting leaf. These manipulations are determined by the rates $k_g$, $k_b$, and $k_s$, the growth rate, branching rate, and switching rate, respectively. Retracting leaves have only a single possible action, retraction, which involves retreating toward the node they are connected to at rate $k_r$.

Degree-one nodes (leaves) are created from branching events, degree-two nodes arise when retracting leaves reach degree-three nodes, and degree-three nodes are created by branching events. The numbers of these node types are given by $N_1$, $N_2$, and $N_3$, respectively, where $N_1 = N_F + N_R$.

Retracting leaves cannot switch back to being a free leaf, though in section 7.1 we discuss an alternative model without this constraint. Retracting leaves keep retracting (continuing through nodes of degree-two) until they reach a node of degree-three, for which it is possible to define an elimination rate, $k_d$. The states and transitions for all nodes and leaves are illustrated in figure S1.

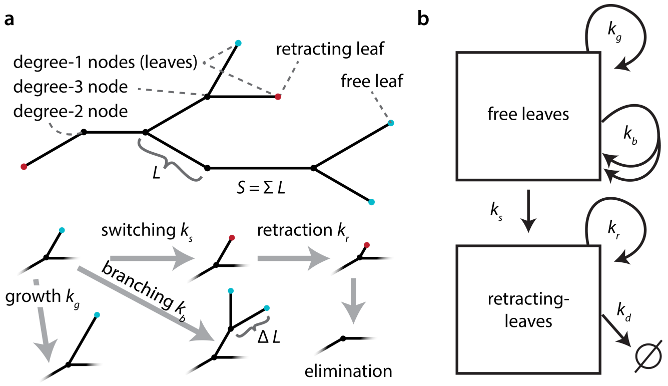

c

| | $k_b$ | $k_g$ | $k_s$ | $k_r$ | $k_d$ |
|---|---|---|---|---|---|
| $S$ | +2 | +1 | 0 | -1 | 0 |
| $N_F$ | +1 | 0 | -1 | 0 | 0 |
| $N_R$ | 0 | 0 | +1 | 0 | -1 |
| $N_1$ | +1 | 0 | 0 | 0 | -1 |
| $N_2$ | 0 | 0 | 0 | 0/-1 | +1 |
| $N_3$ | +1 | 0 | 0 | 0 | -1 |

**Figure S1: Defining restructuring actions. a)** definitions and graphic depictions of restructuring actions. **b)** Possible leaf states and transitions. **c)** Overview of how total size $S$, the number of the two types of leaves $N_F$ and $N_R$, and number the three different node types $N_1$, $N_2$, and $N_3$ change with each transition. Here $\Delta L$ is taken to be 1 for simplicity (Note that the effect of a retraction event on $N_2$ depends on network structure).

Each edge has a length $L_j$ ($j$ being the index of a given edge). We define the network size, $S$, as the sum of all edge lengths, $S = \Sigma L_j$. Branching and growth increase the network size $S$ while retraction decreases the network size (figure S1). Here $\Delta L$ is the length change associated with these transition events, which are defined according to the table in figure S1. The network is not self-avoiding, and edges can overlap without creating new nodes or cycles.

### 4 Network structure

In this section we first derive basic structural properties of this traveling network model such as the edge length distribution, ratios of node types, and relationships between rates using steady state analysis. To understand more complex structural aspects such as the number of node types, we consider a limit case of the full model. Finally we analyze two nondimensionalizations to arrive at a concise description of the model. Our analysis of structure primarily involves analytical results, but we also validate these solutions through numerical simulations of the model.

#### 4.1 Steady state analysis of the model

We first analyze the steady state, in which, despite fluctuations due to stochastic rearrangements, average size $S$ and the average number of free and retracting leaves $N_F$ and $N_R$ are constant over longer time scales.

##### 4.1.1 Basic relationships: $N_F/N_R = 1/R$, $k_s$, and $k_d$

Conservation of size, $S$, leads to (figure S1):

$$\frac{dS}{dt} = 2k_b N_F \Delta L + k_g N_F \Delta L - k_r N_R \Delta L = 0 \implies 2k_b N_F + k_g N_F - k_r N_R = 0. \tag{S1}$$

Similarly, we find for free and retracting leaves in a steady state network:

$$\frac{dN_F}{dt} = k_b N_F - k_s N_F = 0, \tag{S2}$$

and

$$\frac{dN_R}{dt} = k_s N_F - k_d N_R = 0. \tag{S3}$$

From eq. (S2) we find the relationship:

$$k_s = k_b. \tag{S4}$$

The ratio of retracting to free leaves, $R$ can be written from eq. (S1):

$$\frac{N_F}{N_R} = \frac{k_r}{2k_b + k_g} = \frac{1}{R}. \tag{S5}$$

Together with eq. (S3) we find:

$$k_d = k_s \frac{N_F}{N_R} = \frac{k_s}{R}. \tag{S6}$$

From eqs. (S4), (S5), and (S6) then follows:

$$k_d = \frac{k_s k_r}{2k_b + k_g} = \frac{k_b k_r}{2k_b + k_g}. \tag{S7}$$

These arguments provide expressions for $k_s$ (eq. (S4)), $k_d$ (eq. (S7)), and the ratio of free to retracting leaves in terms of $k_b$, $k_g$, and $k_r$. Note that all of these are independent of network size, $S$.

#### 4.2 Average edge length $\bar{L}$ and edge growth time $\tau_g$

Another important feature defining the overall geometry of the network is the average edge length.

##### 4.2.1 Arguments for mean edge length

Consider a free leaf. Upon coming into existence the adjacent edge has length $\Delta L$. This leaf ceases being free with either the next switching or branching event. The growth time, $\tau_g$, is the time it takes for either a branching or switching event, which is the inverse of the sum of switching and branching rates as,

$$\frac{1}{\tau_g} = k_s + k_b = 2k_b. \tag{S8}$$

The last transformation uses the steady state condition eq. (S4). During this time $\tau_g$, the edge will have grown according to $k_g$ to a total length $L_f$ in $\Delta L = 1$ increments. The average length that such an edge grows to before branching or switching is therefore:

$$\bar{L}_f = \Delta L + \tau_g k_g \Delta L, \tag{S9}$$

or with eq. (S8) and $\Delta L = 1$:

$$\bar{L}_f = \Delta L \left(1 + \frac{k_g}{2k_b}\right) = \frac{2k_b + k_g}{2k_b}. \tag{S10}$$

##### 4.2.2 Derivation of edge length distribution

In this section we derive the edge length distribution by writing and analyzing the steady state master equation for the edge lengths considering $\Delta L = 1$ for simplicity. We start with the 'free edges', or edges adjacent to free nodes. Here $L_f$ indicates the length of a free edge and $F_m$ indicates the number of free edges of length $m$. Free edges of length one are created by branching of a free leaf. Growth events increase the length of the associated free edge, and switching events remove the 'free' association from the edge. Figure S2 shows the state diagram and transitions between states.

The steady state equation for $F_1$ is:

$$\frac{dF_1}{dt} = F_1(-k_s - k_b - k_g) + 2k_b \Sigma F = 0 \implies F_1 = \frac{2k_b \Sigma F}{k_s + k_b + k_g}, \tag{S11}$$

where $\Sigma F$ is the sum of all the free edges, $\Sigma F = \sum_{m=1}^{\infty} F_m$.

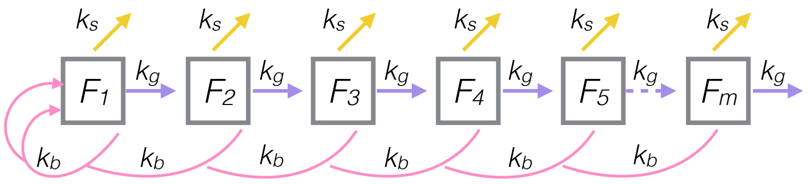

**Figure S2: Steady state visualization of the length of the edges associated with free leaves in the network.** $F_1$ denotes an edge of length $\Delta L = 1$, $F_2$ denotes an edge of length 2, etc. Branching events from nodes associated with edges of any length create 2 new edges of length $\Delta L = 1$ adjacent to free nodes, and cause the initial edge to lose its association with a free node (becoming associated instead with an internal node). Switching events cause the free node to become a retracting node, thus the associated edge loses its free node association, becoming associated instead with an retracting node. Growth events increase the edge length by $\Delta L = 1$.

Similarly we write the steady state equations of $F_2$, $F_3$ and $F_4$ here:

$$\frac{dF_2}{dt} = F_1 k_g - F_2(k_s + k_b + k_g) = 0 \implies F_2 = \frac{k_g F_1}{k_s + k_b + k_g} = k_g \cdot \frac{2k_b \Sigma F}{(k_s + k_b + k_g)^2}, \quad \text{(S12)}$$

$$\frac{dF_3}{dt} = F_2 k_g - F_3(k_s + k_b + k_g) = 0 \implies F_3 = \frac{k_g F_2}{k_s + k_b + k_g} = k_g^2 \cdot \frac{2k_b \Sigma F}{(k_s + k_b + k_g)^3}, \quad \text{(S13)}$$

$$\frac{dF_4}{dt} = F_3 k_g - F_4(k_s + k_b + k_g) = 0 \implies F_4 = \frac{k_g F_3}{k_s + k_b + k_g} = k_g^3 \cdot \frac{2k_b \Sigma F}{(k_s + k_b + k_g)^4}. \quad \text{(S14)}$$

Hence by induction:

$$\frac{dF_m}{dt} = F_{m-1} k_g - F_m(k_s + k_b + k_g) = 0 \implies F_m = \frac{k_g F_{m-1}}{k_s + k_b + k_g} = k_g^{m-1} \cdot \frac{2k_b \Sigma F}{(k_s + k_b + k_g)^m}. \quad \text{(S15)}$$

Thus the probability density function for edges of length $m$ associated with free leaves is:

$$P(L_f = m) = \frac{F_m}{\Sigma F} = k_g^{m-1} \cdot \frac{2k_b}{(k_s + k_b + k_g)^m}. \quad \text{(S16)}$$

The expression for the mean length of such an edge, $\bar{L}_f$, is:

$$\bar{L}_f = \sum_{m=1}^{\infty} m \cdot \frac{F_m}{\Sigma F}. \quad \text{(S17)}$$

Substituting eq. (S16) and simplifying, we find: The mean length, $\bar{L}_f$ is thus

$$\begin{aligned}
\bar{L}_f = &= \sum_{m=1}^{\infty}\left(m \cdot k_g^{m-1}\frac{2k_b}{(k_s+k_b+k_g)^m}\right), \text{ using eq. (S4),} \\
&= 1\cdot\frac{2k_bk_g^0}{2k_b+k_g} + 2\cdot\frac{2k_bk_g^1}{(2k_b+k_g)^2} + 3\cdot\frac{2k_bk_g^2}{(2k_b+k_g)^3} + 4\cdot\frac{2k_bk_g^3}{(2k_b+k_g)^4} + \ldots.. \\
&= \frac{2k_b}{2k_b+k_g}\cdot\left(1 + 2\cdot\left(\frac{k_g}{2k_b+k_g}\right)^1 + 3\cdot\left(\frac{k_g}{2k_b+k_g}\right)^2 + 4\cdot\left(\frac{k_g}{2k_b+k_g}\right)^3 + \ldots.\right) \\
&= \frac{2k_b}{2k_b+k_g}\cdot\left(\sum_{m=1}^{\infty} m\cdot\left(\frac{k_g}{2k_b+k_g}\right)^{m-1}\right) \\
&= \frac{2k_b}{2k_b+k_g}\cdot\frac{(2k_b+k_g)^2}{4k_b^2} \qquad \left(\text{Since } \sum_{m=1}^{\infty} m\cdot x^{m-1} = \frac{1}{(1-x)^2}\right) \\
\bar{L}_f &= \frac{2k_b+k_g}{2k_b} = 1 + \frac{k_g}{2k_b},
\end{aligned} \tag{S18}$$

which is consistent with the intuitive reasoning that led to eq. (S10).

The above probability distribution and average free edge length can also be obtained by recasting the fate of a free leaf as a Bernouli process where 'success' is a either a branching or switching event and 'failure' is a growth event. Success then happens with probability $p = \frac{k_b+k_s}{k_s+k_b+k_g}$. In such a process, the probability to obtain a number of failures, $x$, before success follows a geometric distribution as, $P(x) = (1-p)^x p$. Since the leaves come into existence already with one unit length, $P(L_f = m) = P(x-1) = (1-p)^{x-1}p$, which is consistent with eq. (S16) for which $\bar{L} = 1/p$ which is also consistent with eq. (S10).

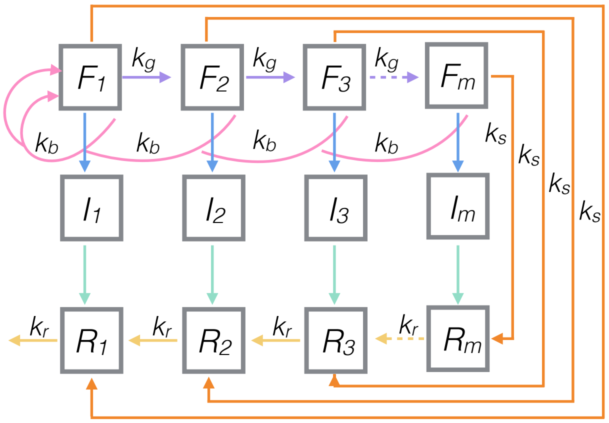

**Figure S3: Steady state visualization of states and transitions between edges associated with different node types.**

Next we calculate the length distribution of edges adjacent to internal nodes. The states and transitions are shown in figure S3 where $I_m$ indicates the number of edges adjacent to internal nodes. Edges associated with internal nodes are eliminated when a retracting leaf adjacent to an edge of length one retracts into a degree-two node ($R_1$ in figure S3). It can be shown that this happens half the time a retracting leaf reaches an internal node (eq. (S42)). The steady state equation for $I_1$ is thus:

$$\frac{dI_1}{dt} = 0 = F_1 k_b - \frac{1}{2} k_r \cdot R_1 \cdot \frac{I_1}{\Sigma I} \implies I_1 = \frac{2F_1 k_b \Sigma I}{R_1 k_r}. \tag{S19}$$

Similarly we write the steady state equations of $I_2$, $I_3$ and $I_4$:

$$\frac{dI_2}{dt} = 0 = F_2 k_b - \frac{1}{2} k_r \cdot R_1 \cdot \frac{I_2}{\Sigma I} \implies I_2 = \frac{2F_2 k_b \Sigma I}{R_1 k_r}, \tag{S20}$$

$$\frac{dI_3}{dt} = 0 = F_3 k_b - \frac{1}{2} k_r \cdot R_1 \cdot \frac{I_3}{\Sigma I} \implies I_3 = \frac{2F_3 k_b \Sigma I}{R_1 k_r}, \tag{S21}$$

$$\frac{dI_4}{dt} = 0 = F_4 k_b - \frac{1}{2} k_r \cdot R_1 \cdot \frac{I_4}{\Sigma I} \implies I_4 = \frac{2F_4 k_b \Sigma I}{R_1 k_r}. \tag{S22}$$

By induction,

$$\frac{dI_m}{dt} = 0 = F_m k_b - \frac{1}{2} k_r \cdot R_1 \cdot \frac{I_m}{\Sigma I} \implies I_m = \frac{2F_m k_b \Sigma I}{R_1 k_r}. \tag{S23}$$

Thus we can write

$$\frac{I_m}{\Sigma I} = \frac{2F_m k_b}{R_1 k_r}. \tag{S24}$$

From the global balance condition (across all edges) in the steady state we can write

$$\Sigma F \cdot 2k_b = R_1 k_r \implies R_1 = \frac{\Sigma F \cdot 2k_b}{k_r}. \tag{S25}$$

Thus by substituting eq. (S25) into eq. (S24) we get

$$\frac{I_m}{\Sigma I} = \frac{2F_m k_b}{R_1 k_r} = \frac{2F_m k_b}{\Sigma F \cdot 2k_b} = \frac{F_m}{\Sigma F}, \tag{S26}$$

which means that the internal edge length distribution matches the free edge length distribution for the network (eq. (S16)). Hence the mean length of these edges associated with internal nodes, $\bar{L}_i$, is also the same as the mean length of edges associated with free leaves,

$$\bar{L}_i = \bar{L}_f. \tag{S27}$$

Eq. (S25) can also be combined eq. (S11) to obtain the equation for length 1 edges associated with retracting nodes:

$$R_1 = \frac{\Sigma F \cdot 2k_b}{k_r} = \frac{\Sigma F \cdot 2k_b}{2k_b + k_g} \cdot \frac{2k_b + k_g}{k_r} = F_1 \cdot \frac{2k_b + k_g}{k_r} \tag{S28}$$

Equations for longer edges (figure S3) associated with retracting leaves can be obtained from the steady state equations:

$$\begin{aligned}
\tfrac{dR_1}{dt} &= 0 = k_rR_2 - k_rR_1 + \frac{1}{2}k_rR_1\frac{I_1}{\Sigma I} + k_sF_1 \\
\implies & -k_rR_2 = -k_rR_1 + \frac{1}{2}k_rR_1\frac{I_1}{\Sigma I} + k_bF_1 \quad (\text{since} \ \ k_s = k_b) \\
\implies & -k_rR_2 = -F_1(2k_b + k_g) + \frac{1}{2}k_rR_1\frac{I_1}{\Sigma I} + k_bF_1 \ (\text{by using eq. (S28)}) \\
\implies & -k_rR_2 = -F_1(2k_b + k_g) + \frac{1}{2}k_rR_1 \cdot \frac{2F_1k_b}{R_1k_r} + k_bF_1 \ (\text{by using eq. (S24)}) \\
\implies & k_rR_2 = F_1(2k_b + k_g) - F_1 \cdot 2k_b \\
\implies & R_2 = F_1\frac{k_g}{k_r} \\
\implies & R_2 = F_2 \cdot \frac{2k_b + k_g}{k_r} \quad (\text{using eq. (S16)})
\end{aligned} \tag{S29}$$

and

$$\begin{aligned}
\tfrac{dR_2}{dt} &= 0 = k_rR_3 - k_rR_2 + \frac{1}{2}k_rR_1\frac{I_2}{\Sigma I} + k_sF_2 \\
\implies & -k_rR_3 = -k_rR_2 + \frac{1}{2}k_rR_1\frac{I_2}{\Sigma I} + k_bF_2 \quad (\text{since} \ \ k_s = k_b) \\
\implies & -k_rR_3 = -F_2(2k_b + k_g) + \frac{1}{2}k_rR_1\frac{I_2}{\Sigma I} + k_bF_2 \ (\text{by using eq. (S29)}) \\
\implies & -k_rR_3 = -F_2(2k_b + k_g) + \frac{1}{2}k_rR_1 \cdot \frac{2F_2k_b}{R_1k_r} + k_bF_2 \ (\text{by using eq. (S24)}) \\
\implies & k_rR_3 = F_2(2k_b + k_g) - F_2 \cdot 2k_b \\
\implies & R_3 = F_2\frac{k_g}{k_r} \\
\implies & R_3 = F_3 \cdot \frac{2k_b + k_g}{k_r} \quad (\text{using eq. (S16)}).
\end{aligned} \tag{S30}$$

By induction

$$R_m = F_m \cdot \frac{2k_b + k_g}{k_r} \tag{S31}$$

is the generalized relation between $R_m$ and $F_m$. The length distribution of edges associated with retracting leaves is the same as edges associated with free leaves and internal nodes. Hence the mean length of edges associated with retracting leaves, $\bar{L}_r$ is also the same as the mean lengths associated with free and internal nodes,

$$\bar{L}_r = \bar{L}_i = \bar{L}_f. \tag{S32}$$

Additionally, the ratio of edges associated with free leaves to edges associated with retracting leaves matches the ratio of free and retracting leaves given in eq. (S5).

Summarizing the findings about edge length, the steady state master equation approach matches the average edge length from intuitive reasoning, and the full edge length distribution of *all* edges - including edges adjacent to free leaves that are still growing, edges adjacent to internal nodes, and edges adjacent to retracting leaves - follows an identical geometric form. Thus rather than considering different average edge lengths we can consider a single average edge length of the network, $\bar{L}$,

$$\bar{L} = \bar{L}_r = \bar{L}_i = \bar{L}_f = \frac{2k_b + k_g}{2k_b}. \tag{S33}$$

Note that here we assumed $\Delta L = 1$, for other $\Delta L$, the following holds,

$$\bar{L} = \Delta L \frac{2k_b + k_g}{2k_b}. \tag{S34}$$

These derived distributions match those extracted from simulations of the model.

#### 4.3 Number of nodes

The number and distribution of different types of nodes are fundamental properties of network structure. This section explores steady state relationships between, and limits for, the numbers of different node types. Section 4.4 develops further insights into the number of nodes by analyzing an informative limit case. The relationships from these two sections allow prediction of a network's structure from its underlying restructuring rates or, conversely, allow systems to set their restructuring rates to achieve desired structures.

##### 4.3.1 Internal nodes, number and elimination: $N_2$, $N_3$, and $k_x$

After having considered the results of imposing a steady state number of leaves (degree-one nodes, i.e., $N_1 = N_F + N_R$) and steady state size, $S$, we next analyze the results of steady state numbers of degree-three nodes and degree-two nodes. Here the size can be divided by the average edge length to obtain the average number of edges, $N_E$, as:

$$N_E = S/\bar{L}. \tag{S35}$$

In general, the following relationship holds for any acyclic connected network with maximum degree of three:

$$N_E = N_1 + N_2 + N_3 - 1. \tag{S36}$$

That eq. (S36) is valid can be seen as a network with a single edge has just two degree-one nodes; any larger network can then be constructed by adding successively a new edge to any of the existing nodes, which adds exactly one edge and one degree-one node while transforming either a degree-one node into a degree-two node, or a degree-two node into a degree-three node; hence for each construction step eq. (S36) is valid.

Additionally, the number of internal degree-three nodes is:

$$N_3 = N_1 - 2. \tag{S37}$$

That eq. (S37) is valid can be show similarly as for eq. (S36): Starting with a single edge, the network has two degree-one nodes and no degree-three nodes. Any additional edge that is attached to an existing degree-one node introduces a new degree-two node, while leaving the number of degree-one nodes and degree-three nodes unchanged; any new edge attached to a degree-two node produces as new degree-three nodes as well as a new degree-one node. Hence any acyclic connected network can be constructed this way, and eq. (S37) is correct for each step. The number of degree-two nodes then follows from eqs. (S36) and (S37):

$$N_2 = N_E + 3 - 2N_1. \tag{S38}$$

Accordingly, from eqs. (S37) and (S38) we also find:

$$N_2 = N_E - 1 - 2N_3. \tag{S39}$$

For the steady state of the degree-three nodes we also find (see figure S1):

$$\frac{dN_3}{dt} = k_b N_F - k_d N_R = 0. \tag{S40}$$

But, eq. (S40) does not provide us with any new insights as it is redundant with combining eqs. (S3) and (S4).

The steady state of the degree-two nodes is given by the balance of new degree-two nodes being generated when a retracting leaf hits a degree-three node (which we already established happens at a rate $k_d$), furthermore by a degree-two node being eliminated when a retracting leaf (i.e., degree-one node) encounters a degree-two node (which happens at a rate which we call $k_x$):

$$\frac{dN_2}{dt} = k_d N_R - k_x N_R = 0. \tag{S41}$$

Which gives

$$k_x = k_d. \tag{S42}$$

Hence in steady state, half of all retraction events that hit a node hit a degree-three node, and the other half hit a degree-two node, so given a node is encountered by a retracting leaf, we can write an average termination probability, $P_t = 1/2$.

Therefore (in steady state) the average length of retraction ($\bar{L}_R$) before eliminating the retracting leaf is:

$$\bar{L}_R = \bar{L}\frac{1}{P_t} = 2\bar{L}. \tag{S43}$$

##### 4.3.2 Bounds on the number of leaves $N_1$, and relationship to $N_F$ and $N_R$

The total number of leaves and leaves of each type (degree-one nodes given by $N_1 = N_F + N_R$) are important quantities since leaves are the sites of active manipulation and restructuring in this model. The ratio of the number of retracting to free leaves is $N_R/N_F = R = \frac{2k_b+k_g}{k_r}$ (see eq. (S5)). Using this ratio and the definition $N_1 = N_F + N_R$ we can express the number free and retracting leaves in terms of $k_g$, $k_b$, $k_r$, and $N_1$ as:

$$N_F = N_1 \frac{k_r}{k_r + 2k_b + k_g}, \tag{S44}$$

$$N_R = N_1 \frac{2k_b + k_g}{k_r + 2k_b + k_g}. \tag{S45}$$

At this point, eqs. (S34) and (S35) allow the number of edges to be determined from the underlying rates $k_b$, $k_g$, $k_r$, and network size $S$. For the three node types, $N_1$, $N_2$, $N_3$ there are two independent equations, eqs. (S36) and (S37). Thus, another independent expression is required to fully resolve the network structure. Such an expression would then also determine $N_F$ and $N_R$ through eqs. (S44) and (S45).

Intuitively, as the size of the network increases, so should the number of each node type. However, finding the form of this relationship turns out to be challenging. For a given size, there are many possible configurations of the network. Enumerating all the possible structures and tabulating their likelihoods is difficult, since these structures emerge and interconvert during the stochastic dynamic restructuring process.

We therefore focus on how the number of leaves, $N_1$, depends on size, $S$, and we first consider two extreme forms of structure: an unbranched chain of nodes and a perfectly branched tree for which we obtain the following relationships:

For an unbranched chain of nodes,

$$N_1 = 2 \text{ (the two ends of the unbranched chain)}, \tag{S46}$$

and for a tree without degree-two nodes,

$$N_1 = \frac{N_E + 3}{2} \text{ (equivalent to a perfectly binary branched tree)}. \tag{S47}$$

Note that these relations follow directly from eq. (S37) with $N_3 = 0$, and eq. (S38) with $N_2 = 0$, respectively.

These extrema set bounds on the largest and smallest number of leaves $N_1$ a network could have at a given size $S$. However, these bounds are too wide to be particularly helpful in gaining deeper understanding of these systems. We next perform a deeper analysis on $N_1$ and its relationship to network structure for a more general and informative set of special cases.

#### 4.4 Network structure in the limit of $k_r \gg k_b$, $\bar{L} = 1$

In this section we analyze the stochastic dynamic branching process that generates the network to determine the network structure including the desired missing relationship between $N_1$ and $S$, the distribution of side branches, and aspects of criticality and self-similarity in the model.

For simplicity, we consider the limit case of rapid retraction ($k_r \gg k_b$). In the case of infinite retraction rate, eq. (S5) shows that all leaves will be free, thus we can simplify the model by ignoring the retracting leaves and edges associated with retracting leaves, hence $N_F = N_1$. We also set $\bar{L} = 1$, eliminating $k_g$ for simplicity. (Note that this last step can be done without loss of generality, shown later in section 4.5.)

Using these simplifications, we consider the possible fates of a newly formed free leaf within a given network under steady state conditions, enumerated in figure S4. For simplicity, the actions of all of the leaves are synchronized and time is discretized by $k_b$, such that at each time step each leaf either 1) branches or 2) switches and fully retracts, with equal probability ($k_b = k_s$), given steady state conditions.

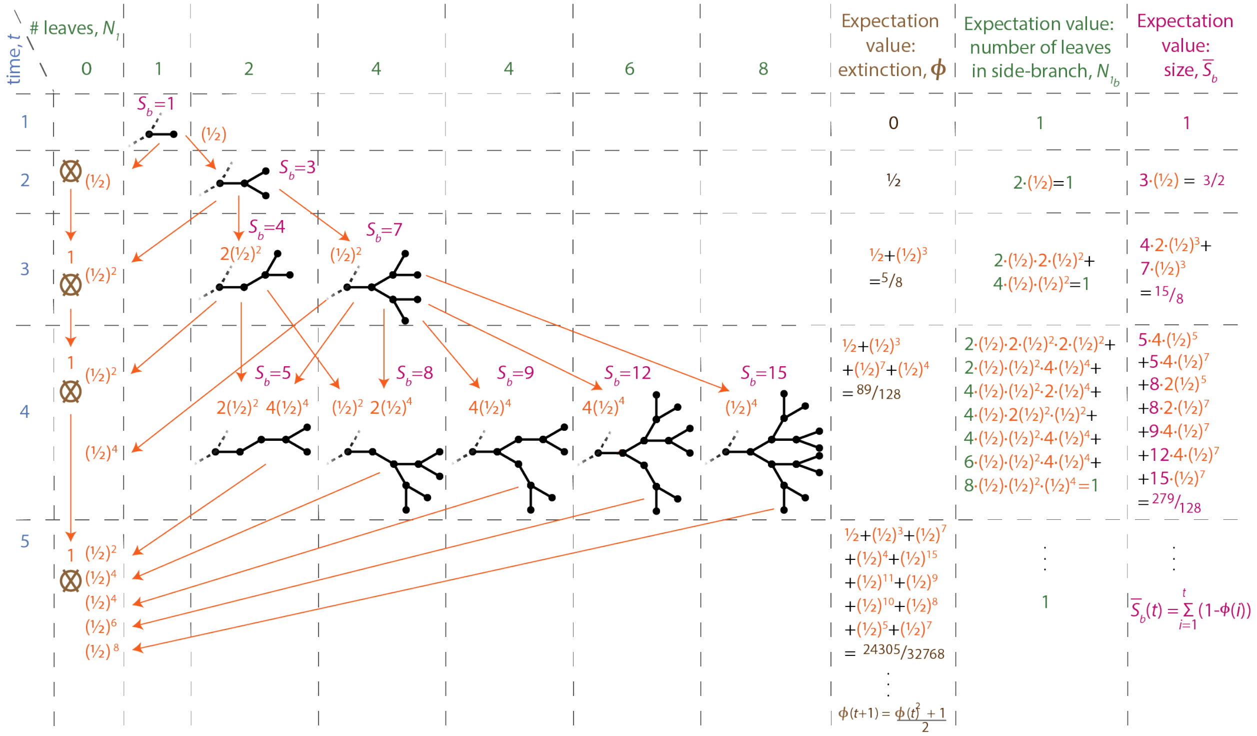

**Figure S4: Transition diagram for a single side branch illustrated through the first four time steps.** The figure displays the possible fates of a side branch assuming infinite retraction speed, no growth, and equally likely synchronized branching or growth for each leaf at each time point. The probability of branch extinction can be written recursively as $\phi(t+1) = \frac{\phi(t)^2+1}{2}$, with $\phi(2) = 1/2$ (OEIS sequence A167424 (*1*)). The expectation value for the number of leaves remains 1 through time. The expectation value for side branch size increases through time as $\bar{S}_b(t) = \sum_{i=1}^{t}(1 - \phi(i))$. Only one structure for each combination of size and leaves is displayed.

A newly formed free leaf may develop into numerous different temporally changing structures and may eventually go extinct. We refer to this entity as a side branch. The possible trajectories of a side branch are illustrated and tabulated exhaustively through the first four time steps in figure S4 (though only one of each representative form is shown). The total number of possible side branch structures rises double exponentially, so displaying structures for further time points becomes cumbersome. Note that we initially assume that these side branches are much smaller than the entire network size, $S$, before considering the case when they are not.

Using this simplified limit case in the following subsections we next compute the expectation values of number of descendent free leaves (4.4.1), probability of extinction (4.4.2), mean side branch size (4.4.3), and ultimately properties of the entire network including: the relationship between the number of edges, number of leaves, and node ages (4.4.4); the distribution of retraction lengths (4.4.7); and the distribution of node-ages (4.4.8). These results are compared to a self-similar approximation (4.4.5) and to simulations that violate the $k_r \gg k_b$ assumption, having $k_r$ and $k_b$ of the same order (4.4.6).

##### 4.4.1 The network is at a critical point

In the steady state we saw earlier that $k_b = k_s$. This implies that the effective branching ratio is 1, i.e, during any given time interval the number of newly generated free leaves is balanced by the number of free leaves switching to the reacted state. If this effective branching ratio were above 1 [or below 1] then the network would be super-critical [or sub-critical] and the number of free leaves would explode [or decay] exponentially – which is not consistent with steady state behavior. The network is thus at a critical point analogous to the critical point in a Galton-Watson process (*2*). This can be seen through the first few time steps of exhaustively enumerated possible fates of a free leaf shown in figure S4, where the expectation value for the number of leaves remains 1, in agreement with the classic result for the mean number of offspring in a critical Galton-Watson process (*2*). We will use this result when later considering the network mean field structure, and in supplementary section 6.2 when we discuss how being at a critical point also enables the network to be an efficient detector of environmental stimuli and to navigate fields of such stimuli. Note that the steady state network remains at this critical point even outside the $k_r \gg k_b$ limit case considered here.

##### 4.4.2 Side branch extinction probability

To understand the side branch fates we first analyze their extinction probability through time. Side branches that fully retract do not start growing again. Thus, extinct side branches are a sink state, and the extinction probability monotonically increases with time. The extinction probability of a side branch over its lifetime can be computed as follows: Let $\phi(t+1|1)$ be the probability that a branch born at time 1 as a single free leaf becomes extinct before or up to $t+1$ time steps. This probability is reported in the third to last column of figure S4. Clearly, $\phi(1|1) = 0$ since $t = 1$ refers to the initial time point when the free leaf has come into existence. At the next time step $\phi(2|1) = 1/2$, which is the relative ratio of switching (and extinction) over all possible scenarios (extinction and branching). (Note: Outside of steady state, this condition would become $\phi(2|1) = k_s/(k_s + k_b)$.)

For later times, the shape of the tree becomes increasingly more complicated and $\phi$ becomes difficult to track, so we will set up a recursion using these initial cases: If the side branch has gone extinct before or up to $t+1$, then it either (i) died in the next time step after birth, which has a probability $\phi(2|1) = 1/2$, or (ii) it branched at the next time step after birth, which has a probability $(1 - \phi(2|1)) = 1/2$, and both of the branches (born at $t = 2$) went extinct before $t+1$. In case (ii), the two branches are independent, hence the probability of *both* dying is the product of probabilities:

$$\phi(t+1|1) = \phi(2|1) + (1 - \phi(2|1)) \cdot \phi(t+1|2) \cdot \phi(t+1|2). \tag{S48}$$

If the process is invariant under time shifts, which is true under steady state (and beyond), then $\phi(t+1|2) = \phi(t|1)$. This just means that each of the 2 new branches, younger by one time step than the original branch, face the same extinction probability as the original branch did one time step earlier. This is true since the same branching and switching probabilities apply to all leaves irrespective of when they arose.

This reasoning leads to the recursion:

$$\phi(t+1|1) = \phi(2|1) + (1-\phi(2|1))\phi(t|1)^2 = \frac{\phi(t|1)^2+1}{2}. \tag{S49}$$

The last equality holds at steady state. In all the following we now drop the initial time for brevity, i.e., $(\cdot) := \phi(\cdot|1)$. This leads to:

$$\phi(t+1) = \frac{\phi(t)^2+1}{2}, \tag{S50}$$

with $t \geq 1$ and $\phi(1) = 0$.

This is equivalent to OEIS sequence A167424 (*1*). A closed form for this expression does not exist, but the recursion can be approximated as,

$$\phi(t) \approx 1 - \frac{2}{t} + \frac{1}{t^2}, \tag{S51}$$

which is more accurate for large $t$, (*3*), or a more exact solution can be obtained from the related OEIS A076628

$$\phi(t) \approx 1 - \frac{2}{t + \ln(t) + 0.767994 + \frac{\ln(t)}{t}} \tag{S52}$$

(Courtesy, Michael Somos and reference (*4*)).

Notice that from eq. (S50) we can derive the recursion for the *survival* probability, $\varphi(t) := 1 - \phi(t)$, as,

$$\varphi(t+1) = \varphi(t) - \frac{\varphi(t)^2}{2}. \tag{S53}$$

The first terms of this formula then correspond to the complement of the extinction probability as illustrated in figure S4:

$$\varphi(2) = \frac{1}{2}$$
$$\varphi(3) = \frac{1}{2} - \frac{1}{2}\cdot\frac{1}{4} = \frac{3}{8}$$

*et cetera.*

The difference equation for successive time steps in eq. (S53) then becomes

$$\Delta\varphi(t) = \varphi(t+1) - \varphi(t) = -\frac{\varphi(t)^2}{2}, \tag{S54}$$

which in a continuum approximation yields the following Cauchy problem:

$$\dot{\varphi}_c(t) = -\frac{\varphi_c^2(t)}{2} \quad \text{with boundary condition} \quad \varphi_c(2) = \frac{1}{2} \tag{S55}$$

which has a unique solution as

$$\phi_c(t) = \frac{1}{1+2/t} \quad \text{and} \tag{S56}$$

$$\varphi_c(t) = \frac{1}{1+t/2}. \tag{S57}$$

These are the solutions for the extinction and survival probabilities of a newly formed side branch (pinned at the first event, at $t = 2$) in a stochastic process that can branch and switch continuously with equal rates rather than with discrete time steps. The survival probability of the continuous process is slightly higher than the discrete one, $\varphi_c(t) \approx \varphi(t)$, (e.g. at $t = 3$, it is $2/5 = 16/40$ instead of $3/8 = 15/40$). Eq. (S56) offers a more concise form than eq. (S52) and is more accurate than eq. (S51), especially at low $t$, hence we use it below.

##### 4.4.3 Side branch size, $S_b$

We next derive the expectation value for the side branch size, $\bar{S}_b(t)$, defined as the average of the sum of all the edge lengths in a side branch at time $t$. Over the first few exhaustively tabulated possibilities in figure S4, we see that $\bar{S}_b(t)$ monotonically increases with $t$, despite the increasing likelihood that the side branch goes completely extinct (given by $\phi(t)$).

Our derivation for the side branch size involves considering edges that arose at different times. At time $t$ a side branch can be composed of edges with ages ranging from 1 to age $t$, which we refer to as different 'age-classes', denoted by index $i$. The expectation value for the size of the entire side branch, $\bar{S}_b$, can be computed by summing the expected size contribution from edges in each age-class. The expected size contribution from edges in each age-class can be computed by multiplying the number of possible edges within each age-class by the probability that they exist. We indicate the number of possible edges in age-class $i$ at time $t$ by $E_i(t)$, and we indicate the probability that an edge in age-class $i$ exists at time $t$ with the function $\theta_i(t)$. Summing this product over all edge age-classes present in the side branch gives the total expected side branch size,

$$\bar{S}_b(t) = \sum_{i=1}^{t} E_i(t)\theta_i(t). \tag{S58}$$

Finding expressions for $E_i(t)$ and $\theta_i(t)$ will thus enable computation of the side branch size.

Starting with the expression for $E_i(t)$, only one edge can be as old as the side branch (age-class $i = t$). This is the 'founding edge' that was present when the side branch was created. It is possible to have more numerous edges of younger ages (figure S4 and figure S5). It is straightforward to see that the number of possible edges in each age-class for a side branch of total age $t$ doubles with decreasing age-class as,

$$E_i(t) = 2^{t-i}, \tag{S59}$$

where $1 \leq i \leq t$.

Focusing next on the expression for $\theta_i(t)$, the probability of existing for edges in each age-class can be computed by considering the rules that built the side branch. We first consider the youngest age-class, then the oldest, then generalize by considering intermediate age-classes.

Regardless of total side branch age, $t$, the youngest edges in age-class $i = 1$ must have a probability of existing of 1 divided by the number of possible edges in the age-class, $\theta_1(t) = \frac{1}{E_1(t)=2^{t-1}}$, to preserve the critical expectation value of 1 leaf. This age-class always contributes $\theta_1(t) * E_1(t) = \frac{2^{t-1}}{2^{t-1}} = 1$ to the expected side branch size.

The existence probability of the innermost 'founding edge' (age-class $i = t$) is the same as the probability that the entire side branch exists at all, $\theta_{i=t}(t) = 1 - \phi(t)$ (as derived in 4.4.2), since this edge exists if and only if the side branch exists. This age-class then contributes $\theta_{i=t}(t) * E_t(t) = (1 - \phi(t)) * 1$ to the expected side branch size.

The existence probability for edges in intermediate age-classes can be computed by beginning with the critical expectation of one leaf at each time step and following the subsequent recursive process that builds the structure. Starting with a side branch of arbitrary age $t$, select an age-class $i \geq 2$. The edges in this age-class arose at an earlier time, $t_e = t - i$, and there are $E_i(t) = 2^{t-i} = 2^{t_e}$ possible edges in this age-class (by eq. (S59)). Upon birth at time $t_e$, these edges must have had an initial existence probability, $\theta_i(t_e)$, of $\theta_i(t_e) = \frac{1}{2^{t_e}}$ following the same arguments above to preserve the critical average expectation of one leaf at time $t_e$. Subsequently, each of these edges experienced the same recursive rules that govern the single 'founding edge' in a side branch, which has the temporally changing probability of existence, $(1 - \phi(t))$ (as derived in 4.4.2). This means that at time $t_e + 1$ the existence probability of edges in the age-class under consideration is $\theta_i(t_e+1) = \theta_i(t_e)*(1-\phi(2)) = \frac{1}{2^{t_e}}*(1-\phi(2))$, or their initial probability of existence at time $t_e$ multiplied by the chance that they exist after another time step. Continuing forward to time $t$, this becomes $\theta_i(t) = \frac{1}{2^{t_e}} * (1 - \phi(t - t_e))$.

By substituting $t-i$ for $t_e$ we find that the existence probability for edges in each age-class in a side branch of age $t$ is given by,

$$\theta_i(t) = \frac{1 - \phi(i)}{2^{t-i}}. \tag{S60}$$

Note that this expression is also valid for the the oldest and youngest age-classes.

Having found expressions for $E_i(t)$ and $\theta_i(t)$, we can substitute eqs. (S59) and (S60) into eq. (S58) to get the expected size of a side branch of age $t$ as,

$$\bar{S}_b(t) = \sum_{i=1}^{t}(1 - \phi(i)) = \sum_{i=1}^{t} \varphi(i), \tag{S61}$$

which is consistent with the tabulated sizes in figure S4.

###### 4.4.4 Size of the entire network, $S$, and relationship to leaves, $N_1$

We now shift from considering the fate of a single 'founding leaf' giving rise to a side branch of small size to considering the entire network. The entire network at any given time emerged at some point in the past from a single free leaf according to the same sort of stochastic process that gave rise to each side branch. It might then seem possible to use the results directly from the idealized scenario for side branches to make inferences about the entire network's structure. However, there is a key difference between a side branch and the entire network that must first be addressed. Individual side branches may arise and go entirely extinct, but for most systems the whole network maintains a roughly constant size, $S$, and is not permitted to retract completely and go extinct. Exactly calculating the network properties requires the full set of states and transition probabilities which depend on how network existence is enforced and how size is conserved. In our simulations, for example, we maintain the network size by multiple conditions: requiring $k_s = k_b$ to ensure steady state, dependence of the switching rate on a hill function involving the network size, and an additional constraint that the last two free leaves never switch (see section 8.1.1 for more detail on the implementation). Hence in most systems when the side branch sizes approach the network size, the assumption that they can go entirely extinct becomes poor.

It would still be desirable to utilize the results above on the key properties of side branches derived from detailed considerations of the recursive stochastic process. In order to utilize these results, yet account for the condition that the network is enforced to exist we modify the scenario depicted in figure S4 by eliminating the transition to complete extinction. The conceptual picture is as follows. Consider a network of size $S$ at an arbitrary time. This network arose from a single founding leaf at an earlier time. The same recursive stochastic rules described above for side branches governed the development of the structure that emerged from this founding leaf with the key distinction that the branch never went extinct. Instead, this structure continued developing until it 'consumed' all the network's size and became the entire network. To approximately capture the fact that the entire network is not permitted go extinct we modify the considerations for small side branches for the whole network by eliminating the transition to the zero-leaf 'extinct' sink state depicted in figure S4 (second column from the left, labeled '# leaves $N_1 = 0$'). In the progression of advancing from one time step to the next, this is essentially equivalent to considering the same process for generating side branches above where the specific outcome of complete extinction is canceled by rerolling the dice should all leaves stochastically retract simultaneously during any given time step.

The weighted average calculations (right columns of figure S4) for the expected number of leaves and the expected size can then be modified accordingly for the absence of the sink state. The sink state contributes zero leaves and zero size, and its probability increases over time according to eq. (S50). The new expectation values for the entire network, conditioned by the absence of the sink state, can be computed by dividing the side branch expectation values by the survival probability, $\varphi(t) := 1 - \phi(t)$ (see eq. (S53)).

The expectation value for the number of leaves of the entire network over time conditioned in this way is no longer '1' as it was for each side branch, but now instead increases monotonically as:

$$\bar{N}_{1_{net}}(t) = (\bar{N}_{1_b}(t)| \neq \emptyset) = \frac{\bar{N}_{1_b}(t)}{\varphi(t)} = \frac{1}{\varphi(t)}, \tag{S62}$$

where the '$net$' and '$b$' subscripts have been added to emphasize which quantities are for the entire network and which are for small side branches.

The conditioned size of the network increases as,

$$\bar{S}_{net}(t) = (\bar{S}_b(t)| \neq \emptyset) = \frac{\bar{S}_b(t)}{\varphi(t)} = \frac{\sum_{i=1}^{t} \varphi(i)}{\varphi(t)}, \tag{S63}$$

where again the '*net*' and '*b*' subscripts have been added to emphasize which quantities are for the entire network and which are for small side branches. The probabilistic appearance of one such network is depicted schematically in figure S5.

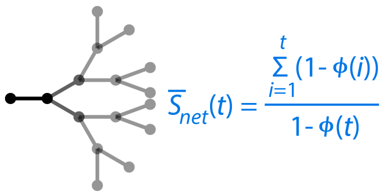

**Figure S5: Size of the network.** The network size after a time $t$ is computed by multiplying the number of edges of each age by the probability of their existence. This size is then modified according to the condition that the network exists (see eq. (S63)). Opacity indicates the probability of existence for each edge, averaged over all possibilities, schematically shown here for an oldest node of age $t = 4$.

Equation (S63) allows us to predict the typical age of the oldest node for a network of a given size. Equation (S62) allows us predict the typical number of leaves that the network has at a given age. Combining these two equations allows us to relate the typical number of leaves to the typical size. Obtaining a single closed form equation without the recursive form of $\varphi(t)$ or sum in eq. (S63) would be helpful to more clearly illuminate the relationship between the number of leaves and size. In the following we make various simplifications to arrive at such an approximate closed form.

We first substitute the approximate form for $\varphi(t)$ from eq. (S57). Equation (S62) then becomes, dropping the '*net*' subscript and bar notation,

$$N_1(t) \approx 1 + t/2. \tag{S64}$$

Equation (S63) then becomes, dropping the '*net*' subscript and bar notation,

$$S(t) \approx \left( \sum_{i=1}^{t} \frac{1}{1 + i/2} \right) (1 + t/2), \tag{S65}$$

which can be written as,

$$S(t) \approx \frac{1}{2}(t + 2)(2H_{t+2} - 3) \tag{S66}$$

where $H_i$ is the ith harmonic number ($H_i = \sum_{j=1}^{i} \frac{1}{j}$). The harmonic numbers can be approximated as, $H_t \approx \ln(t) + \gamma + ...$, where $\gamma$ is the Euler–Mascheroni constant, $\gamma = 0.577216...$, resulting in a closed form approximation for the relationship between network size and age,

$$S(t) \approx \frac{1}{2}(t + 2)(2(\ln(t + 2) + \gamma) - 3). \tag{S67}$$

Despite the approximations, eq. (S67) and eq. (S64) do a good job capturing the full recursive sum from eq. (S63) and the recursion from eq. (S62), with $< 10\%$ error for $t > 20$ as shown in figure S6.

Finally, substitution of eq. (S64) into eq. (S67) relates the number of leaves and network size,

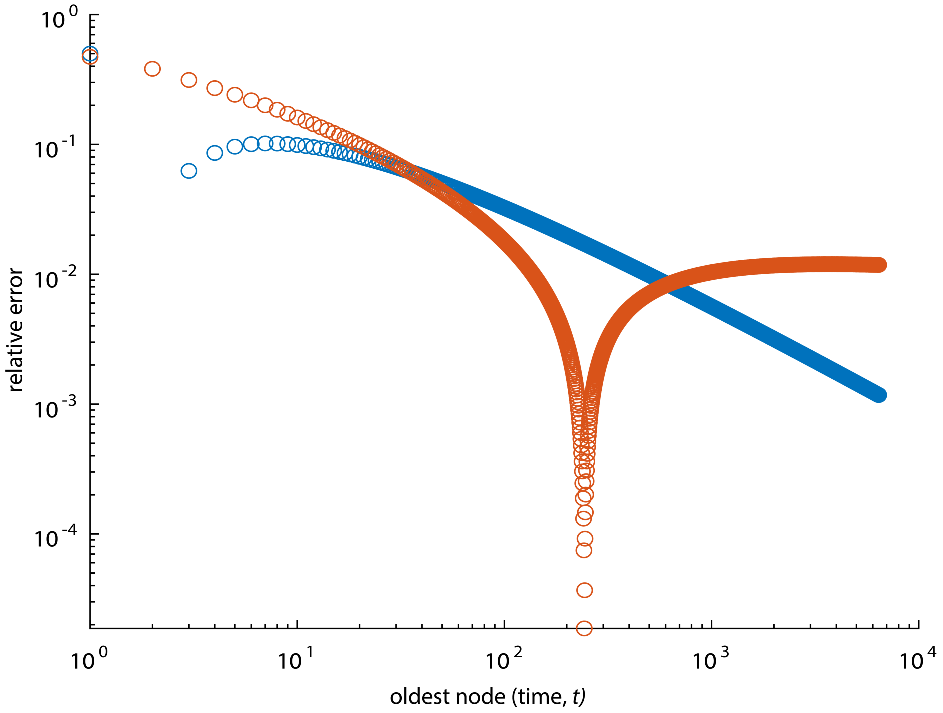

**Figure S6: Relative error of closed form approximations.** Relative error was calculated for network size as $\left|\frac{(\frac{1}{2}(t+2)(2(\ln(t+2)+\gamma)-3))-\left(\sum_{i=1}^{t}\varphi(i)\right)/\varphi(t)}{\left(\sum_{i=1}^{t}\varphi(i)\right)/\varphi(t)}\right|$ (red) and number of leaves as $\left|\frac{(1+t/2)-1/\varphi(t)}{1/\varphi(t)}\right|$ (blue) to compare the full iteratively calculated solutions (eqs. (S63) and (S62)) to closed forms (eqs. (S67) and (S64)). The sharp dip in relative error of the size around $t = 230$ is due to the approximation transitioning from less than to greater than the exact form, the difference passing through zero.

$$\bar{S} \approx N_1(2\ln(2N_1) + c), \tag{S68}$$

where $c = 2\gamma - 3 = -1.845568...$ Which can also be written as,

$$S \approx N_1(2\ln(N_1) + c_1), \tag{S69}$$

where $c_1 = 2\ln(2) + 2\gamma - 3 = -0.46...$

This closed form approximation provides a relationship between the two key network properties of size $S$ and number of leaves $N_1$.

###### 4.4.5 Self-similar approximate mean field structure of the network

Next we show how an intuitive heuristic approach can be used to generate an approximate mean field structure and arrive at a similar relationship between $N_E$ and $N_1$ as the analysis above in sections 4.4.2-4.4.4.

We consider the network generating process described at the beginning of section 4.4, $k_r \gg k_b$, $\bar{L} = 1$. This allows us to simplify the model by eliminating growth and accounting for leaves only in the free state. One could derive an analytical relationship between $N_E$ and $N_1$ for any given $S = N_E$ by averaging over all possible network configurations and their likelihoods, but doing so is not straightforward as seen in the previous section. Instead, we generate an approximate explicit structure representative of the mean field to capture essential properties. Such a representative structure should possess the following key attributes:

1) age equivalence - nodes of the same age should be interchangeable. Note that this also establishes symmetry at each branching point.

2) preserve criticality - each leaf should have, on average, one descendent leaf, as established in section 4.4.1.

A structure such as the one in figure S7 possesses both of these attributes. With time discretized according to $k_b$ and leaves synchronized such that each leaf underwent one branching or switching event per time point, as discussed in section 4.4, each leaf is connected by a single unique path to the oldest node of age $a_{max}$. Along this path, for example the one highlighted in blue in figure S7, bifurcations occur with degree-three nodes. We refer to each bifurcation as a 'level', where $\ell$ indicates the level, from 0 at the leaves up to a maximum level $\mathcal{L}$. Due to age equivalence, when proceeding along the path from youngest to oldest node the number of offspring leaves doubles with increasing level with each such bifurcation encountered. To maintain an average critical expectation of one offspring leaf per side branch, a corresponding set of degree-two nodes (having side branches with zero offspring leaves) must be inserted to average out the leaves encountered at the bifurcated degree-three node.

This results in a self-similar structure that is more densely branched near the leaves and more sparse with increasing age. Such a self-similar structure can also be generated by scaling the structure by a factor of 2 at each increasing level, or by considering a perfectly branched binary tree and removing all descendant nodes from nodes of all ages that are not a power of 2, i.e. keeping the descendants from nodes with ages 1,2,4,8,...$2^i$ (figure S7).

This structure allows us to establish a relationship between the number of leaves and

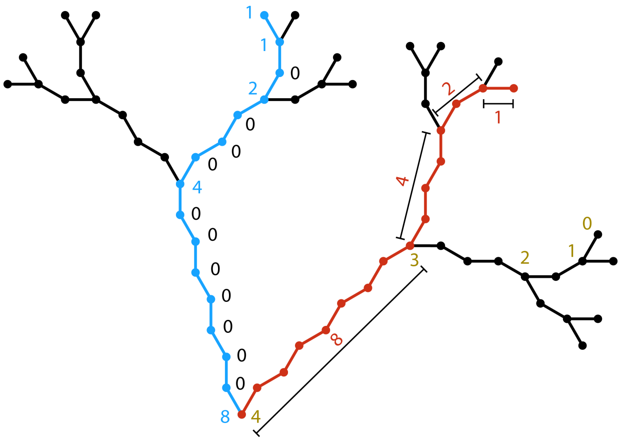

| Max Level, $\mathcal{L}$ | New leaves | Cummulative leaves, $N_1$ | Size, $S$ | New length along path | Cummulative length along path, $a_{max}$ |
|---|---|---|---|---|---|
| 0 | - | 1 | 0 | - | 0 |
| 1 | 1 | 2 | 2 | 1 | 1 |
| 2 | 2 | 4 | 8 | 2 | 3 |
| 3 | 4 | 8 | 24 | 4 | 7 |
| 4 | 8 | 16 | 64 | 8 | 15 |
| $\mathcal{L}$ | $2^{\mathcal{L}-1}$ | $2^{\mathcal{L}}$ | $\mathcal{L}2^{\mathcal{L}}$ | $2^{\mathcal{L}-1}$ | $2^{\mathcal{L}}$-1 |

**Figure S7: Mean field form of the network.** A representative explicit mean field structure satisfying age equivalence of nodes and the critical expectation of leaf offspring. The table indicates how network attributes change with increasing level, color coded as in the structure

the total network size. Various attributes of the structure at different levels are given in the table in figure S7. Of interest here, the total number of leaves increases with maximum level $\mathcal{L}$ as,

$$N_1(\mathcal{L}) = 1 + \sum_{\ell=1}^{\mathcal{L}} 2^{\ell-1} = 2^{\mathcal{L}}. \tag{S70}$$

The total size increases as,

$$N_E(\mathcal{L}) = \sum_{\ell=1}^{\mathcal{L}} \frac{1}{\ell} \cdot \ell 2^{\mathcal{L}} = \mathcal{L}2^{\mathcal{L}}. \tag{S71}$$

The maximum level, $\mathcal{L}$, can be written as $\mathcal{L} = \log_2(N_1)$ and substituted into the expression for size to give an expression relating size and number of leaves,

$$N_E = S = N_1 \log_2(N_1) = N_1 \ln(N_1)/\ln(2). \tag{S72}$$

Hence we recover the $S \approx N_1 \ln(N_1)$ relationship of eq. (S69) but with different correction factors.

This structure also allows us to compute the age $a_{max}$ of a typical 'oldest node' in the network by counting the edges along the path from old to young nodes (cumulative length along the path in figure S7). Since this path is created at rate $2k_b$ (sections 5.1 and 5.3), the cumulative length along the path divided by the rate the path is created gives the age of the oldest node in real time,

$$a_{max} = 2^{\mathcal{L}} - 1 = \frac{N_1 - 1}{2k_b} \sim \frac{N_1}{2k_b}. \tag{S73}$$

Note that branching angle and whether the branching direction went right or left will change the appearance of the network, but have no impact on the aspects under consideration in this section.

###### 4.4.6 Analytical predictions of size, $S$, and leaves, $N_1$ vs simulations

Eq. (S68) and eq. (S72) relate $N_E(= S)$ and $N_1$ (given in sections 4.4.4 and 4.4.5) allowing us to solve the expressions in sections 4.3.1 and 4.3.2 for any and all the node types as a function of $N_E$. This provides a useful description of the network structure. As we we will see in the following sections, it allows us to derive a relocation time, understand the characteristic network widths, and ultimately rationalize high level network search behaviors. We note that the form of the equations relating $N_E$ and $N_1$ is, perhaps not coincidentally, reminiscent of the form for the mean path length in a randomly increasing binary tree (*5*).

To derive these expressions we assumed that retraction was fast relative to branching ($k_r \gg k_b$). In the full model, this is not always the case, and we assess this assumption by comparing the derived expressions to data from the full simulation in figure S8. We find reasonable agreement between the predicted and simulated numbers of leaves over a few orders of magnitude of effective network size. The self-similar heuristic approach leading to eq. (S72) tends to overestimate the number of leaves by about 25%, and tends to perform worse for larger networks with more edges (figure S8a). The recursive solution and its approximation also overestimate the number of simulated free leaves but matches the full simulations better than the self-similar prediction (figure S8, also seen in the main text figure 2). For both predictions the discrepancy is only mildly worse with increasing $K_{B^*}$ (figure S8a), $K_{B^*} = \frac{k_b \bar{L}}{k_r \Delta L}$, discussed in section 4.5.2) despite violation of the $k_r \gg k_b$ assumption at higher $K_{B^*}$. This might be understood as some network size being 'trapped' in actively retracting branches that were not included in the simplified reasoning. The remaining discrepancy for the self-similar approximation does not appear to be due to violating the $k_r \gg k_b$ assumption since the discrepancy remains at very small $K_{B^*}$ where this assumption is valid. For small $K_{B^*}$, for both predictions, the discrepancy with simulations is mildly worse for larger network sizes (figure S8b).

###### 4.4.7 Retraction length distribution in the limit of $k_r \gg k_b$, $\bar{L} = 1$

Systems operating at critical points often display self-similar characteristics. Beyond the self-similar structure heuristically predicted in 4.4.5, we next describe how the model can be recast as a simple set of algorithmic rules and analyze the self-similar distribution of retraction events.

Here we again consider the limit case of $k_r \gg k_b$, $\bar{L} = 1$, now to understand the distribution in the number of $\bar{L}$ edges eliminated by retracting nodes before a degree-three node is reached, which we refer to as a 'retraction length', $L_R$. Note that this is different from the distribution of edge lengths themselves, which is covered in section 4.2 (here edge lengths are set to 1, $\bar{L} = 1$). This limit case can be captured by recasting the model as a simple set of dynamical rules governing how such a network changes over time:

i) Begin with a binary tree.

ii) If the total network size exceeds the threshold size, $S$, then randomly select a leaf and fully retract it past all degree-two nodes until a degree-three node is reached or only two nodes remain.

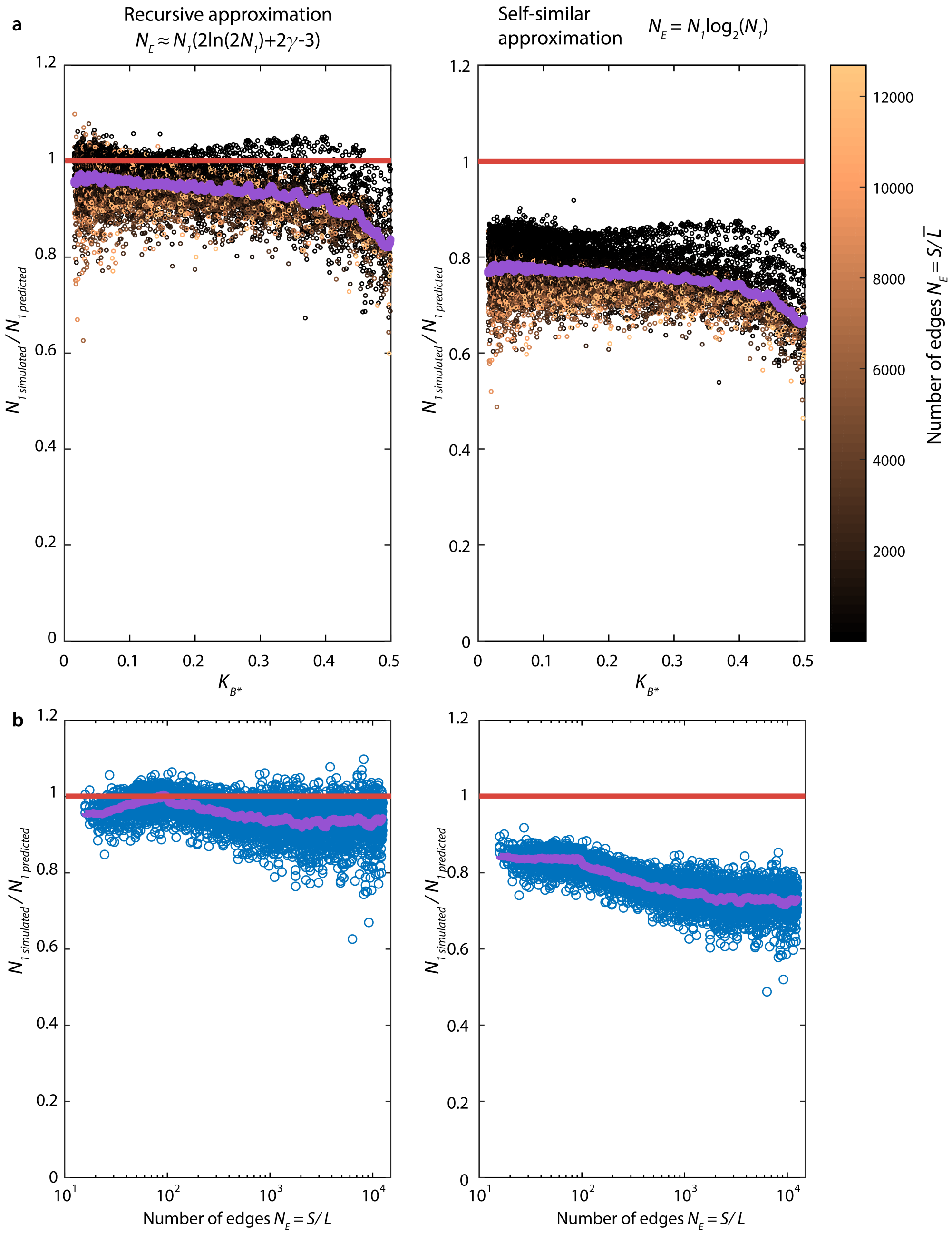

**Figure S8: Analytical predictions for $N_1$ capture simulations over three orders of magnitude of network size. a)** The recursive approximation matches the simulation results better than the self-similar approximation, though both overestimate the number of leaves compared to results from the stochastic simulation. Both predictions are modestly worse at larger $K_{B^*}$, where the assumption of fast retraction is poorest. Color indicates the number of edges in the network. The mauve line indicates a moving average. **b)** For $K_{B^*} < 0.25$, both predictions vary across network sizes in their ability to predict the number of leaves in the stochastic simulation. The recursive approximation is fairly consistent in performance across network sizes. The self-similar approximation matches the stochastic simulation best at small network sizes and performs worst for larger networks. The mauve line indicates a moving average.

iii) Randomly select a leaf and branch it.

iv) Return to ii.

Under these rules, the retraction happens completely before the next branching event (consistent with $k_r \gg k_b$), thus branching events can be seen as slow driving events, and retraction events can be seen as rapid relaxations. This framing of the model helps to highlight the relationship of this limit case to self-organized criticality (SOC) (*6*). In SOC systems, rapid relaxations (classically 'avalanches') can be scale free with probability distributions that obey power laws (*7*). If this is also the case for retraction events in this model, the probability, $P(L_R)$, of a retraction event of length $L_R$ should then follow a form of the type:

$$P(L_R) = cL_R^a \tag{S74}$$

where $c$ and $a$ are constants. This should sum to one such that,

$$1 = c\sum_{L_R=1}^{\infty} L_R^a. \tag{S75}$$

The mean retraction length can be given by the sum of the probability of each retraction length multiplied by that length as

$$\bar{L}_R = c\sum_{L_R=1}^{\infty} L_R \cdot L_R^a. \tag{S76}$$

We know this mean retraction length to be 2 from above (eq. (S43)) such that

$$2 = c\sum_{L_R=1}^{\infty} L_R \cdot L_R^a. \tag{S77}$$

Combining the mean (eq. (S77)) and probability distributions (eq. (S75)) we find that

$$\frac{1}{2} = \frac{c\sum\limits_{L_R=1}^{\infty} L_R^a}{c\sum\limits_{L_R=1}^{\infty} L_R \cdot L_R^a} \tag{S78}$$

which can be written in terms of the Riemann Zeta function, $\zeta$ as

$$\frac{1}{2} = \frac{\zeta(-a)}{\zeta(-a-1)} \tag{S79}$$

and can be solved to give $a \approx -2.47875....$ By substituting into eq. (S75) or eq. (S77), $c \approx 0.7408....$

$$P(L_R) = cL_R^a \tag{S80}$$

$$c = 0.7408...$$

$$a = -2.47875...$$

This result shows good agreement with the simulation for probabilities of retraction events of different sizes in this simplified model (figure S9), demonstrating how constants can be derived for emergent distributions displayed by the model.

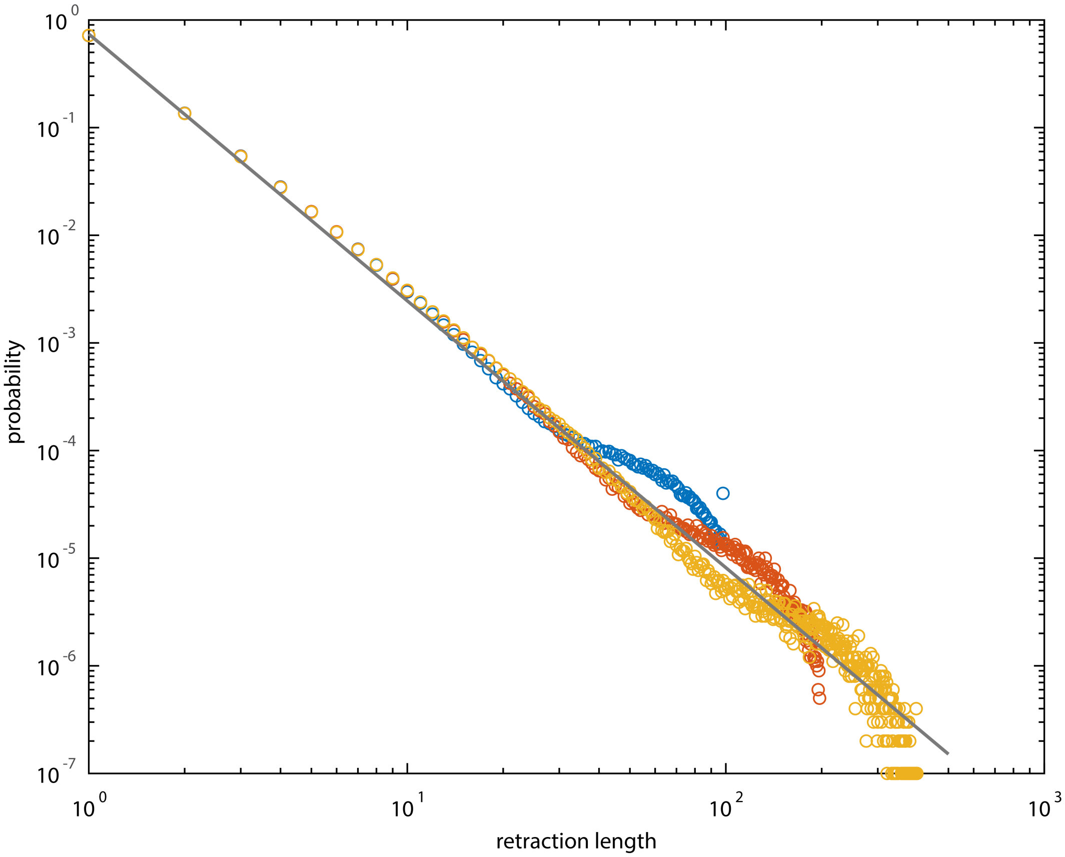

**Figure S9: Probability distribution of retraction event lengths is scale free.** Gray is the analytical power law prediction (eq. (S80)), points are data from simulated networks of sizes $S$ =100 (blue), 200 (red), and 400 (gold) over $10^7$ branching events. Networks simulated as described in section 8.2.

###### 4.4.8 Self-similarity of the network, node age distribution

We note that, qualitatively, within the network structure the most abundant nodes are the youngest, and older parts of the network are ‘thinner’, giving the network a fractal-like appearance (see supplementary movie 2 and main paper figure 2a). Self-similar distributions are often exhibited by systems at critical points, and the heuristic arguments in section 4.4.5 predict a self-similar structure. Considering the ‘age-class’, $i$, of nodes as in section 4.4.3, the heuristic arguments in section 4.4.5 predict a power law probability distribution with exponent -1 for nodes of different age-classes, $P(i) \propto i^{-1}$. We next adapt the arguments from section 4.4.1-4.4.4 to derive the node-age probability distribution and to better understand the network structure and self-similar properties.

The probability that a node has a given age within the network is the typical number of nodes in that age-class, $N_i$, divided by the total number of nodes in the network,

$$P(i) = \frac{N_i}{N_{total}}. \tag{S81}$$

For a binary tree, the total number of nodes is $N_t = N_E + 1$ (eq. (S36)). Neglecting the slight difference of '1', we follow the arguments in section 4.4.3 concerning edges. The number of nodes of each age-class can be written as the product of the total number of nodes possible in that age class and their probability of existence within all possible node of that age. For a network of age $t$, there are $2^{t-i}$ possible nodes in each age class (see eq. (S59)), and following from eq. (S60) for side branches, and the arguments in section 4.4.4 conditioning the probability on the network's existence, the probability of each possible node in a class is

$$\theta_{i_{net}}(t) = (\theta_{i_b}(t) | \neq \emptyset) = \frac{\frac{\varphi(i)}{2^{t-i}}}{\varphi(t)} \tag{S82}$$

where the '$net$' and '$b$' subscripts have been added to emphasize which quantities are for the entire network and which are for small side branches.

The expected number of nodes in each age-class is then,

$$N_i = \frac{2^{t-i} \frac{\varphi(i)}{2^{t-i}}}{\varphi(t)}. \tag{S83}$$

Under the $\bar{L} = 1$ assumption, the total number of nodes is essentially the network size, $S \approx N_{total}$ (eq. (S36)), which leads after substitution along with a simplified eq. (S83) into eq. (S81) to,

$$P(i) = \frac{\varphi(i)}{S\varphi(t)}. \tag{S84}$$

Recognizing that $1/\varphi(t)$ is in fact simply the number of leaves, $N_1$ (see eq. (S62)), and substituting eq. (S57), eq. (S84) can also be expressed as,

$$P(i) = \frac{1}{1 + i/2} \frac{N_1}{S}. \tag{S85}$$

Using eq. (S68), $N_1$ can be written in terms of $S$. Substituting this for $N_1$ in eq. (S85) gives the probability distribution for node age within a network of a given size.

The results of this prediction are plotted with simulations of three different network sizes in figure S10a. A 'finite age' effect occurs around a maximum age for each network size, but otherwise the analytical arguments capture the distributions from simulation, indicating these arguments are indeed capturing key essential features of the network structure.

We note that in eq. (S85) as $i \to \infty$ (older nodes which are found in larger networks), $P(i) \to c/i$, matching the -1 power law form predicted by the simple self-similar arguments in section 4.4.5. Compared to perfect self-similarity, the youngest nodes are slightly less common (figure S10b) which is likely the cause of the discrepancy between the predictions in section 4.4.5 and the simulation results for the relationship between size and the number of leaves (figure S8).

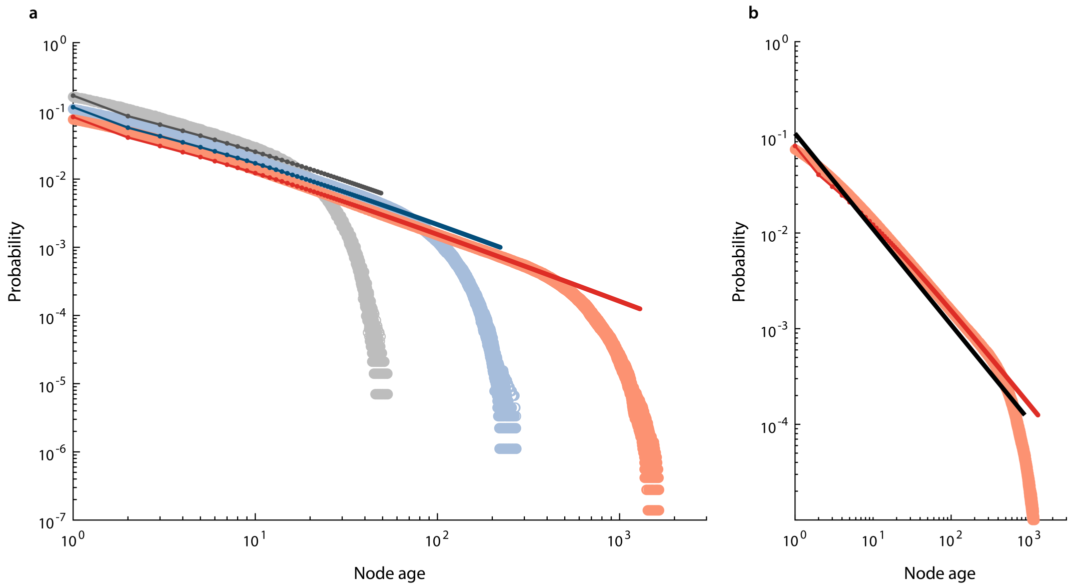

**Figure S10: Node age distribution.** **a)** Node age distributions for networks of sizes $S = 160$ (gray), 1000 (blue), and 8000 (red). Darker lines in the foreground are the analytical prediction (eq. (S84)). Lighter points in the background are from simulations. **b)** Comparison between the analytical predicted distribution and simulated distribution for $S = 8000$ (same as in panel a) with the exact power law distribution predicted by perfect self-similarity (solid black line, see section 4.4.5). Networks simulated as described in section 8.1 with $K_B = 0.01$, $K_G = 0$.

#### 4.5 Nondimensionalization

Note that of the six rates discussed above $k_b$, $k_g$, $k_s$, $k_r$, $k_d$, and $k_x$, in the steady state $k_d$ and $k_x$ are emergent and $k_s = k_b$ so only three are truly independent. The time scale and length scale in the system can be chosen to further collapse the model. Here we analyze two nondimensionalized versions of the model, first nondimensionalizing time by the retraction time to explicitly preserve the impacts of growth and branching rates, then nondimensionalizing both time and length to understand a compact universal form of the model.

##### 4.5.1 Nondimensionalization by retraction time

Here we use $k_r$ to nondimensionalize the time scale, leading to redefinition of $K_R := k_r/k_r = 1$, $K_B := k_b/k_r$, $K_G := k_g/k_r$, $K_S := k_s/k_r$, and $K_D := k_d/k_r$. This results in key steady state relationships of:

$$K_S = K_B$$

$$\frac{N_R}{N_F} = R = 2K_B + K_G$$

$$K_D = K_X = \frac{K_B}{2K_B+K_G}$$

$$\bar{L} = \Delta L \cdot \frac{2K_B+K_G}{2K_B}$$

$$\bar{L}_R = 2\bar{L}$$

$f = 2K_B + K_G$ (see section 5.3)

$2K_B + K_G \leq 1$ (see section 5.3)

$D^{\bullet} := \frac{D}{k_r} = \frac{\Delta L^2 (2K_B + K_G)^2}{8K_B} \cdot \frac{1+\cos(\alpha/2)}{1-\cos(\alpha/2)}$ (in two dimensions, see section 5.2)

$\tau_r^{\bullet} := \tau_r k_r = \frac{a_{max}}{2K_B} \approx \frac{N_1}{2K_B}$ (see section 5.4)

Note that each of the above relationships are only dependent on the rescaled branching and growth rates, $K_B$ and $K_G$ (additionally branching angle for the diffusivity and size for the relocation time). Nondimensionalization by retraction time is the normalization primarily discussed in the main text.

###### 4.5.2 Nondimensionalizing time scale and length scale

Here we use $\frac{\bar{L}}{k_r \Delta L}$ (the time scale for a retracting leaf to reach the next node) to nondimensionalize the time scale and $\bar{L}$ to nondimensionalize the length scale, leading to: $K_{R^*} := \frac{k_r \bar{L}}{k_r \Delta L} = \frac{\bar{L}}{\Delta L}$, $K_{B^*} := \frac{k_b \bar{L}}{k_r \Delta L} = \frac{2k_b + k_g}{2k_r}$, $K_{G^*} := \frac{k_g \bar{L}}{k_r \Delta L}$, $K_{S^*} := \frac{k_s \bar{L}}{k_r \Delta L}$, and $K_{D^*} := \frac{k_d \bar{L}}{k_r \Delta L}$ and key steady state relationships of:

$K_{S^*} = K_{B^*}$

$\frac{N_R}{N_F} = R = 2K_{B^*}$

$K_{D^*} = K_{X^*} = 1/2$

$\bar{L}^* = \frac{\bar{L}}{\bar{L}} = 1$

$\bar{L}_R^* := \frac{\bar{L}_R}{\bar{L}} = 2$

$f = 2K_{B^*}$ (see section 5.3)

$K_{B^*} \leq 1/2$ (see section 5.3)

$D^* := D \cdot \frac{\bar{L}}{k_r \Delta L} \cdot \frac{1}{\bar{L}^2} = \frac{K_{B^*}}{2} \cdot \frac{1+\cos(\alpha/2)}{1-\cos(\alpha/2)}$ (in two dimensions, see section 5.2)

$\tau_r^* := \tau_r \frac{k_r \Delta L}{\bar{L}} = \frac{a_{max}}{2K_{B^*}} \approx \frac{N_1}{2K_{B^*}}$ (see section 5.4)

Though the diffusivity depends on branching angle and relocation time depends on the number of network edges, most of the key attributes can be expressed solely in terms of the rescaled branching rate, $K_{B^*}$. $K_{B^*}$ can be interpreted as the number of branching events occurring for each free leaf during the time it takes the retracting leaves to reach the next node. Relationships between nodes of different degrees, $N_1$, $N_2$, and $N_3$, are graph properties that are largely independent of the restructuring rates or any normalization, with the exception of a weak dependence on $K_{B^*}$. The next section shows how $K_{B^*}$ can be used to capture a concise universal form of the model.

#### 4.6 Reduced parameter space and universal form

Section 4.5.2 showed that many rescaled network features depend only on $K_{B^*}$, and section 4.4 showed that node numbers depend primarily on $N_E$, weakly on $K_{B^*}$, and on nothing else. Thus any two networks that share these two parameters belong to the same family and have the same rescaled features (rescaled as in section 4.5.2), despite having potentially different underlying unscaled restructuring rates. It is then possible to construct a compact 'universal form' of the model for each family (figure S11) with no growth ($k'_g = 0$), edges of uniform length ($\Delta L' = \bar{L}' = 1$), and retraction rate of one ($k'_r = 1$); (parameters of the universal form are denoted by a 'prime' symbol). This universal form acts as a concise simplified model that shares all key rescaled attributes with other members of its family.

**a**

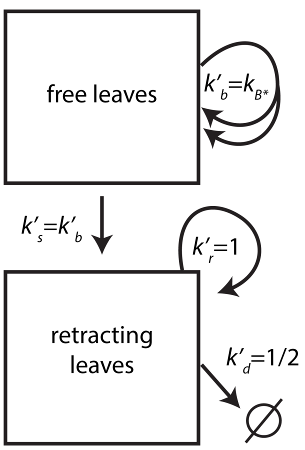

**b**

| | $k'_b$ | $k'_s$ | $k'_r$ | $k'_d$ |
|---|---|---|---|---|
| $S$ | +2 | 0 | -1 | 0 |
| $N_F$ | +1 | -1 | 0 | 0 |
| $N_R$ | 0 | +1 | 0 | -1 |
| $N_1$ | +1 | 0 | 0 | -1 |
| $N_2$ | 0 | 0 | 0/-1 | +1 |
| $N_3$ | +1 | 0 | 0 | -1 |

**Figure S11: Universal form of the model. a)** Transitions between states. Prime symbols denote rates in the universal form. **b)** Table of effects of different restructuring options on node types and size.

In order for any two networks to be in the same family, they must have the same $K_{B^*}$ and $N_E$, thus when constructing the universal form for a given family, $K'_{B^*} = K_{B^*}$. In the universal form, $k'_g = 0$ and $k'_r = 1$, and the remaining free rate, $k'_b$ must be set to give the desired $K'_{B^*}$. We know that, $K'_{B^*} = \frac{2k'_b + k'_g}{2k'_r}$, and the convenient choice of $k'_g = 0$ and $k'_r = 1$ in the universal model allows substitution that gives $k'_b = K'_{B^*}$, hence $k'_b = K_{B^*}$. Thus, to map any model with any combination of $k_b$, $k_g$, and $k_r$ to its corresponding compact universal form, only $K_{B^*}$ needs to be known from the original model and this value can be used directly for the branching rate $k'_b$ in the universal form.

Sections 4.1-4.4 discuss the structure of the network, which depends on the additional parameter, $S$, the network size. Apart from a weak effect of $K_{B^*}$ on the number of degree-one nodes (figure S8), the number of nodes of different degrees depends only on the number of edges, $N_E = S/\bar{L}$. In the universal form above $S' = N'_E$, since $\bar{L}' = 1$. With respect to the number of nodes of different types, any network can then be mapped onto a corresponding universal form by setting $S' = S/\bar{L} = N_E$. Edge lengths in this universal form are uniform, whereas in the full model they follow a geometric distribution. If accounting for this variation is considered important for a certain application, then the edge lengths in the universal form could be drawn from the geometric distribution of the type derived in section 4.2, eq. (S16),

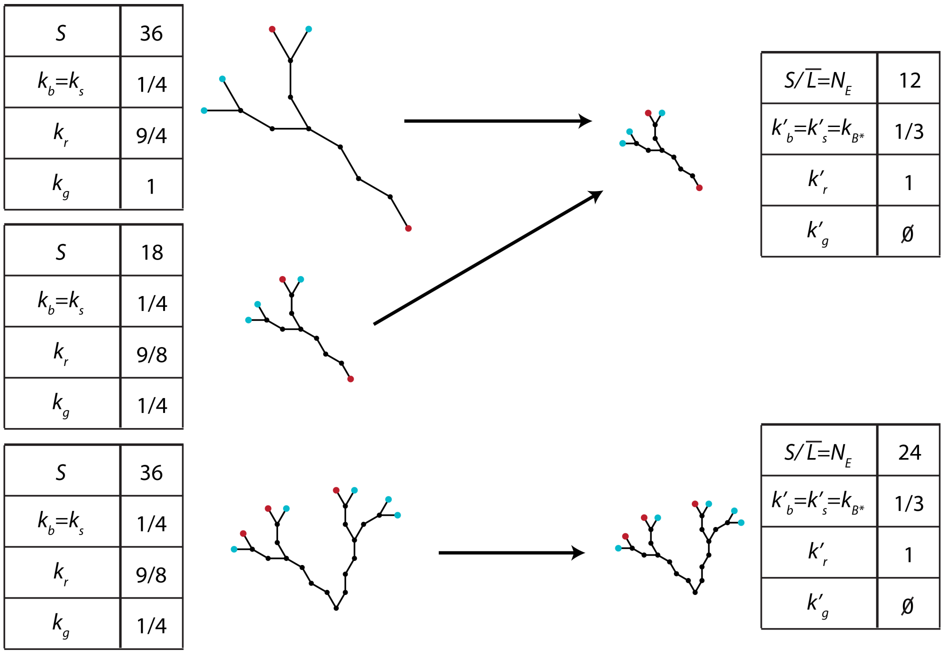

**Figure S12: Illustration showing three example cases rescaled to their corresponding universal forms.**

without changing the mean behavior.

Thus, networks with any combination of $k_b$, $k_g$, $k_r$, and $S$ can be mapped to their corresponding universal forms (figure S12), and these forms can be understood in a 2D parameter space of $K_{B^*}$ and $N_E$ (figure S13). Given the dependence of the rates on each other and the dependence of the numbers of nodes of different degrees on each other, other parameters combinations could also be chosen to perform these normalizations, and the choice of normalization depends on objective.

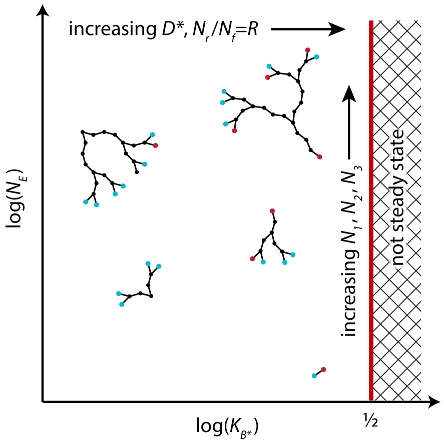

**Figure S13: Key properties of structure and movement in the fully rescaled model depend on $N_E$ and $K_{B^*}$.**

Note that the branching angle is an additional parameter in the model. This parameter influences the spatial embedding of the networks but has no impact on lengths, emergent rates, velocity of leaves, node-type distributions, or connective arrangement of the nodes. Tuning of the branching angle is, however, important for the appearance of the network, and the network movement through a specific space as shown in the next section.

### 5 Network movement

With an understanding of the network's structure, we next sought to understand the movement of the network.

#### 5.1 Movement of the network's head and tail along a path

Consider a network that is spatially embedded with some initial position. Dynamic restructuring causes the network to move, and, after a sufficient amount of time, none of the network's original nodes will remain and the network will occupy a new position. For an acyclic model such as this one, a single unique path can be traced in retrospect between the initial and final positions.

It is informative to identify and consider two special nodes of interest, the 'tail' and the 'head' (see main paper figure 2). The tail is the oldest node in the network. It follows the path on the lagging end of the network. The tail alternates between being a retracting leaf moving along the path at speed $\Delta L \cdot k_r$, and a stationary degree-two node. We term the fraction of time the tail spends moving as $f$, which is not specified but arises automatically from the network's structure and dynamics. The head leads the network, growing and branching along the path. Without any biasing, each free leaf has equal probability of becoming the head, thus the head can only be defined in retrospect. Supplementary movie 4 explicitly illustrates the path, tail, and head concepts.

The head dynamics can be understood by considering a size balance of the head and tail. The head adds size to the network at rate $\Delta L(2k_b + k_g)$ while the tail removes size from the network along the path at rate $f \cdot \Delta L \cdot k_r$. In steady state, the size added by the head must balance with the size removed by the tail, since all other branching events that depart from the path eventually retract back to the path and have no net impact on the steady state size of the network, so,

$$\Delta L(2k_b + k_g) = f \cdot \Delta L \cdot k_r \implies 2k_b + k_g = f \cdot k_r. \tag{S86}$$

Understanding this size balance also helps us to understand the average speed of the head, $v_{head}$, and the average speed of the tail, $v_{tail}$. In steady state, the head and tail must move with the same average speed, $v_{head} = v_{tail}$, otherwise the tail would overtake the head or the head would outpace the tail and the network length along the path would grow without bound. The average speed of the tail is equal to the rate that the tail removes size from the network,

$$v_{tail} = f \cdot \Delta L \cdot k_r. \tag{S87}$$

To balance this, it follows from eq. (S86) that the average speed of the head must be

$$v_{head} = f \cdot \Delta L \cdot k_r = \Delta L \cdot (2k_b + k_g). \tag{S88}$$

Note that $v_{head}$ is also the characteristic speed at which the entire network moves along the path, since in steady state the network size is constant.

#### 5.2 Long time scale diffusive behavior

Of particular interest is understanding how this traveling network model moves through space over longer time scales. Since the network follows the head along the path, understanding the motion of the head allows us to understand the shape of the path and, over long time scales, the average motion of the entire network.

The head moves stochastically along the path with speed $\Delta L \cdot (2k_b + k_g)$ (eq. (S88)), branching at rate $2k_b$ and growing at rate $k_g$. This motion is a correlated random walk of the same type analyzed in the freely rotating chain model in classical polymer physics (*8*). From polymer theory, the end-to-end vector, $\boldsymbol{R}$, is a quantity of interest. The mean squared end-to-end distance of polymer molecules whose backbones are made up of links is:

$$\langle \boldsymbol{R}^2 \rangle = N_{links} l^2 \frac{1 + \cos(\theta)}{1 - \cos(\theta)}, \tag{S89}$$

where $\theta$ is the angle of rotation between links, $l$ is link length, and $N_{links}$ is the number of links. This equation is valid for large $N_{links}$; the derivations and more detail can be found in Doi and Edwards 1988 (*8*) and in Gedde 2013 (*9*). Note that the trivial case of $\theta = 0$ is ballistic, not diffusive.

Links in the polymer backbone correspond to steps in the random walk process we consider here. The number of steps in the random walk is given by the branching rate of the head multiplied by the runtime, $N_{links} \implies \Delta t \cdot 2k_b$. Half the branching angle is equivalent to the angle of rotation $\theta \implies \alpha/2$, and the link length corresponds to the average edge length, $l \implies \bar{L}$, allowing us to write the mean squared displacement of the head, $MSD_{head}$ as

$$MSD_{head} = \Delta t \cdot 2k_b \bar{L}^2 \frac{1 + \cos(\alpha/2)}{1 - \cos(\alpha/2)} \tag{S90}$$

using eq. (S34),

$$MSD_{head} = \Delta t \cdot \frac{\Delta L^2 (2k_b + k_g)^2}{2k_b} \frac{1 + \cos(\alpha/2)}{1 - \cos(\alpha/2)}. \tag{S91}$$

In general, the diffusivity, $D$, can be computed as

$$D = \frac{MSD}{q \Delta t}. \tag{S92}$$

In two dimensions, the constant $q = 4$, allowing us to write $D$ in terms of the underlying restructuring rates and branching angle:

$$D = \frac{\Delta L^2 (2k_b + k_g)^2}{8k_b} \frac{1 + \cos(\alpha/2)}{1 - \cos(\alpha/2)}. \tag{S93}$$

From numerical simulations (see main paper figure 2g), we find that the center of mass of these traveling networks moves superdiffusively at short time scale and transitions to diffusive behavior on longer time scales with the diffusivity matching eq. (S93). Notice that the size of the network does not enter into this expression, indicating that over long time scales the network motion does not depend on network size. From numerical simulations, larger networks take longer to transition from superdiffusive to diffusive behavior, but reach the same diffusivity as smaller networks (figure S14).

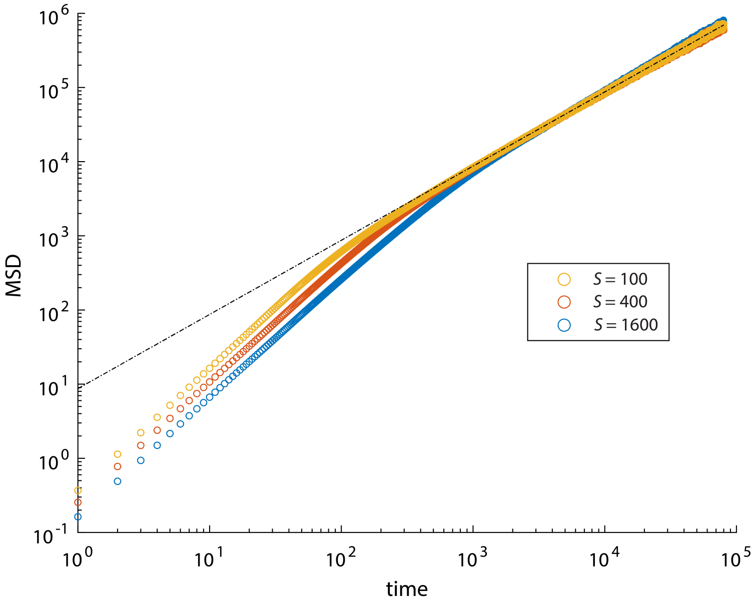

**Figure S14: Mean squared displacement vs time for three different network sizes.** Larger networks tend to take longer to transition to diffusive behavior. Network parameters $k_b = 0.2$, $k_g = 0.1$, $\alpha = 60$ over $\Delta t = 16000000$ shown here. The dashed black line is the analytical diffusive prediction.

##### 5.2.1 Persistence length of the network path

The same understanding that the path is a correlated random walk allows us to again borrow from polymer theory (*10*) and define a persistence length, $L_p$, (the characteristic length scale over which the random walk loses directional correlation) for the path, written in terms of the underlying restructuring rates and branching angle:

$$L_p = -\bar{L}\frac{1}{\ln(\cos(\alpha/2))} = -\frac{\Delta L(2k_b + k_g)}{2k_b \ln(\cos(\alpha/2))}. \tag{S94}$$

#### 5.3 Limits to rates in the steady state

The head, tail, and path concepts and relationships discussed in section 5.1 also establish limits to the allowable parameter combinations in steady state. In eq. (S86), $f$, the fraction of time the tail is moving, cannot be larger than 1, such that,

$$2k_b + k_g \leq k_r \text{ or when normalized by retraction time, } 2K_B + K_G \leq 1. \tag{S95}$$

This limit is consistent with numerical simulations in which the network grows without bound when $2k_b + k_g \geq k_r$.

#### 5.4 Time scale of relocation

We define the 'relocation time scale', $\tau_r$, as the typical time after which none of the original nodes remain. This is a fundamental quantity that distinguishes traveling networks from other types of networks, such as networks that expand from a fixed rooted position, or networks that restructure but never leave the original set of nodes.

The relocation time scale is nearly equivalent to (i.e., slightly longer than) the age of the oldest node in the network, which can be obtained from the arguments about the network structure in section 4.4. Here we provide an approximate estimate for $\tau_r$ by considering the mean-field structure given in section 4.4.5 (figure S7) and the process constructing it. In this structure, the newest nodes are the freshly branched leaves at level 0 and these are connected to the oldest node at the highest level, $\mathcal{L}$. The cumulative number of edges along the path is equivalent to the maximum node age, $a_{max}$, according to eq. (S73), $a_{max} = 2^{\mathcal{L}} - 1 = \frac{N_1 - 1}{2k_b} \sim \frac{N_1}{2k_b}$ (figure S7). This establishes the maximum age within the time discretized by $k_b$ as described in section 4. To compute $\tau_r$ in units of real time, this age must be divided by $2k_b$ resulting in,

$$\tau_r = \frac{a_{max}}{2k_b} \approx \frac{N_1}{2k_b}. \tag{S96}$$

Simulated networks agree reasonably with this prediction (figure S15), with a linear increase in the mean age of the oldest node with $\frac{N_1}{2k_b}$, ($N_1$ computed from network size by eq. (S72)), but with a $\sim 25\%$ offset from the analytical prediction. We note that a significant finite age effect occurs for these networks which impacts the probability distribution of the oldest nodes (figure S10), such that these arguments concerning the relocation time are best considered as an order of magnitude estimate.

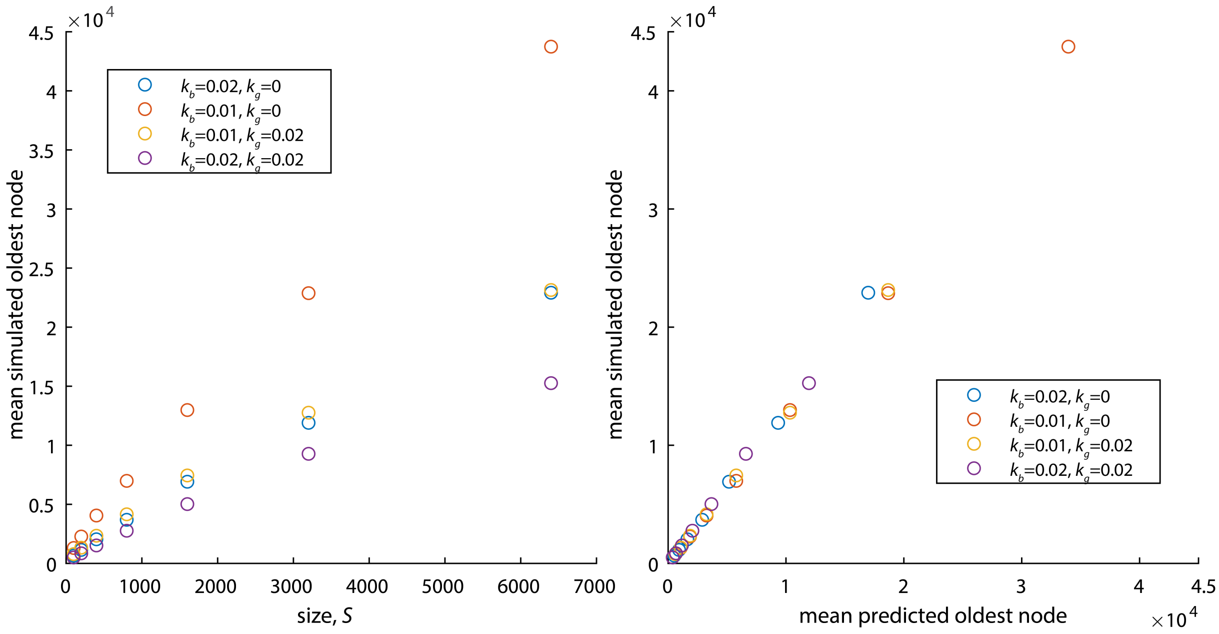

**Figure S15: Average oldest node in simulations matches analytical prediction. a)** Average oldest node vs. network size. **b)** simulated mean oldest node vs. analytically predicted mean oldest node. Age of oldest node is predicted by $\frac{N_1}{2k_b}$. Unity represents perfect agreement. Network parameters for both plots: (blue: $k_b = 0.02$, $k_g = 0$), (red: $k_b = 0.01$, $k_g = 0$), (gold: $k_b = 0.01$, $k_g = 0.02$) (purple: $k_b = 0.02$, $k_g = 0.02$) shown here. $N_1$ computed from network size by eq. (S72)

### 6 Network search

Search is a common reason to travel, and our focus on how networks travel through space allows us to explore their efficacy at various search tasks. Section 5 presented some notion of search by analyzing the long-term network diffusion. Here we investigate additional features of search, first considering unbiased search from stochastic restructuring, then considering search when network restructuring and network movement are biased by the network's environment.

#### 6.1 Unbiased, steady state spatial search

In the following we analyze how the network searches space through unbiased stochastic restructuring. We consider the extent to which the network spreads around the correlated random walk of the path and the granularity of area visited to understand how such networks might set their restructuring rates to optimize for various aspects of search.

##### 6.1.1 Typical width of the network along the path

In section 5 we derived how the head moves as a correlated random walk, superdiffusively over short time scales and diffusively over long time scales. During this motion, the network's various side branches (main paper figure 2f, supplementary movie 4) explore some characteristic distance away from the path. We call this typical search distance away from the path the characteristic network width, $w$.

Understanding the typical network width requires knowledge of how the typical structure of the network depends on underlying restructuring rates and size, particularly the length and shape of the branches departing from the path. The lengths and configurations of these side branches are stochastic, and since an exhaustive probabilistic landscape of network structures is challenging to catalog and consider, here we assume a network structure of the approximate mean field-type discussed in section 4.4.5. This allows us to write the width, $w$ in terms of the underlying restructuring rates, branching angle, and network size.

Considering the mean field structure in section 4.4.5, each route from oldest to youngest node is equally likely to become the path. Thus each of these routes has the same correlated random walk structure as discussed in section 5, with persistence length, $L_p$, (eq. (S94)) and diffusivity, $D$, (eq. (S93)). However, only one route will actually become the path and the other routes can be identified retrospectively as 'side branches'. Whichever route is eventually the path, there remain side branches with chains of up to $a_{max}$ edges departing from the path, where $a_{max} = 2^{\mathcal{L}} - 1$ (eq. (S73)). This is the 'cumulative length along the path' in figure S7, and closely related to the relocation time (section 5.4). The number of edges given by $a_{max}$ can be related to the total number of edges and network size as $N_E = S/\bar{L} = (a_{max} + 1) \log_2(a_{max} + 1)$. These side branches then have a typical maximum length $L_{SB} = \bar{L} a_{max}$. To determine the width of exploration away from the path, it is informative to consider two extreme possibilities for how the the persistence length relates to the side branch length, $L_p$ vs $L_{SB}$:

(Regime 1) If the persistence length is much greater than the side branch length, $L_p >> L_{SB}$, then the side branches can roughly be considered as rods that stick out from the path where $\alpha$ determines how far they reach out perpendicular from the path. We then obtain using simple trigonometry a width of exploration away from the path of,

$$w \sim L_{SB} \sin(\alpha) = \bar{L} a_{max} \sin(\alpha), \tag{S97}$$

(see for example, main paper figure 3d, $\alpha = 15°$).

(Regime 2) In the other extreme, if the persistence length is much less than the side branch length, $L_p << L_{SB}$, the side branches essentially diffuse, and, following the arguments in section 5, the characteristic length scale that these side branches travel away from the path is:

$$w \sim \sqrt{MSD}/2 \sim \sqrt{\frac{a_{max}}{4} \bar{L}^2 \frac{1 + \cos(\alpha/2)}{1 - \cos(\alpha/2)}}, \tag{S98}$$

with the divisor of two to account for distance only in the direction orthogonal to the path (see for example, main paper figure 3d, $\alpha = 90°$).

These two regimes have different implications for search. In the first case when $L_p >> L_{SB}$ the network searches an area surrounding the path of width $w$ in a 'swept area' manner (see schematic in main paper figure 3c). Networks in this regime travel over a wide range, since with a large persistence length they also tend to diffuse rapidly, but these networks do not search thoroughly. In the second case when $L_p << L_{SB}$, the side branches diffuse, and since the persistence length is the same for the side branches and the path, the side branches routinely cross themselves and the path. In this regime, the network performs a thorough, fine scale, and highly redundant search, but the small persistence length keeps this search within a very local area. When $L_p \sim w$, as in main paper figure 3d, $\alpha = 45°$, the network is somewhere between these two extremes and does a search that is intermediate in exhaustiveness, redundancy, and scope.

Note that in the comparison of $L_{SB}$ and $L_p$, only the side branch length, $L_{SB}$, depends on the network size, $S$, so for otherwise equivalent attributes larger networks are more likely to be in the second regime. Only the persistence length, $L_p$ depends on branching angle, $\alpha$. The side branch length and persistence length both depend on the restructuring rates $k_b$ and $k_g$ through $\bar{L}$. Increasing $\bar{L}$ increases both of these lengths, but increases the persistence length more than the side branch length. Hence, for networks of a given network size, there are multiple ways to modulate network search behavior; decreasing $\alpha$ (main paper figure 3d) or increasing $\bar{L}$ are effective ways to transition to the first regime.

##### 6.1.2 Rate of linear exploration by the network

Another important quantity for search is how many new points are covered per unit time by the network. Length is added to the network at free leaves, so new length is proportional to the number of free leaves and the rate that they add length. For each free leaf, length is added at a rate of $\Delta L(2k_b + k_g)$ (see also eq. (S1)), or, new sections of length $\bar{L}$ are added at a rate $2k_b$, such that the new length, $L_{new}$, is added at rate,

$$L_{new} = N_F \Delta L(2k_b + k_g) = N_F 2k_b \bar{L}. \tag{S99}$$

Maximizing $k_b$ increases the addition of new length in three ways, 1) increasing $L_{new}$ directly through the $k_b$ term in eq. (S99), 2) increasing $N_F$ by increasing the fraction of leaves in the free state (see eq. (S5)), and (3) increasing the number of free leaves by decreasing $\bar{L}$, maximizing edges, and maximizing the total number of leaves (see eqs. (S33), (S35), and (S72)).

##### 6.1.3 Area covered by the network

The area covered by a network over time gives a metric for how much space it was able to search. However, the overlaid network shapes are irregular and made up of 1-dimensional lines (main paper figure 3), prompting us to use the box counting method from fractal or Hausdorff dimension to quantify the area covered (*11*). This involves tiling boxes of different sizes over the historical network trajectories and counting the number of boxes with at least one line crossing them. When the number of boxes covered is plotted against the box size in a log-log plot, a regular polygon, such as a rectangle, gives a slope or Hausdorff dimension of -2, and a single line gives a slope or Hausdorff dimension of -1. Overlaid network histories from this model quantified with box counting have typical slopes between -1 and -2, though are not linear over all length scales (figure S16). Details of the implementation can be found in supplementary section 8.3.

This method of quantifying how much area is covered provides a metric for search efficacy at different resolutions. For a given search resolution, a better searcher will collect more boxes of the resolution size. Both the slopes, intercepts, and non-linear features from box counting plots depend on the restructuring rates, as seen in main paper figure 3b. Data from main paper figure 3b show that depending on box size, networks with different restructuring rates collect the most boxes, hence the optimal restructuring rates depend on the search resolution.

Here we explore how the branching angle and duration of search impact search efficacy at different resolutions. With the exception of a few simple arguments and limit cases, these investigations are mostly numerical. Full derivation of the complex structures of these self-crossing, stochastic networks is beyond the scope of this work.

##### 6.1.4 Change in dimension with time

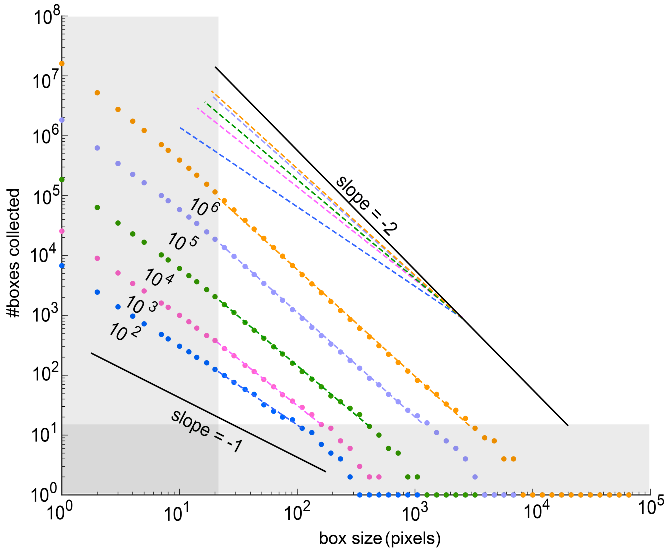

**Figure S16: Hausdorf dimension of the searched area changes over time.** Networks were simulated over different runtimes ($10^2$ to $10^6$ as indicated in the figure), and box counting was used to quantify the area covered at different resolutions and determine the dimensionality of the space searched. Slope of -2 indicates 2D coverage and slope of -1 indicates a 1D coverage. The search dimension tends to progress upward in dimensionality over time. The gray regions were not included when estimating the slope. The left gray region is beneath the pixel length of $\bar{L}$ and the bottom gray region has 10 or fewer counts and is thus highly susceptible to noise. Dashed lines indicating slope are offset and stacked to better compare. Network parameters: $\alpha = 60°$, $S = 100$, $k_b = 0.01$, $k_g = 0$, $k_r = 1$.

A Brownian trail in 2 or more dimensions can be shown to have a Hausdorff dimension of 2 over long times (*11*). As discussed in section 5.2, the path follows a correlated random walk. To better understand how the Hausdorff dimension of these traveling networks changes over time and how it compares to the classical result from Brownian motion we ran simulations for several orders of magnitude of runtime. We kept all other factors constant and performed box counting (*11*) (supplementary section 8.3). We found that the system progresses from a dimensionality of slightly greater than 1 monotonically upward over time, perhaps approaching the Brownian result (figure S16). The increase in dimensionality is likely due to the network crossing over its previous positions and 'filling in' the gaps resulting in a more filled 2D search history. As time progresses, this self-crossing becomes more frequent, which also means that per time, search becomes more redundant and less effective at finding new area, consistent with the sublinear increase in boxes collected with runtime (figure S17).

Since networks of any persistence length eventually move diffusively (except for the trivial case of $\alpha = 0$), networks across the entire parameter space will likely follow this trend of increasing search dimension and increasing search redundancy, though the rate of increase of the dimensionality likely depends on how long it takes to transition from the superdiffusive to diffusive regime. We investigate one such case below with networks having different branching angles.

##### 6.1.5 Effect of branching angle, $\alpha$, on search over time

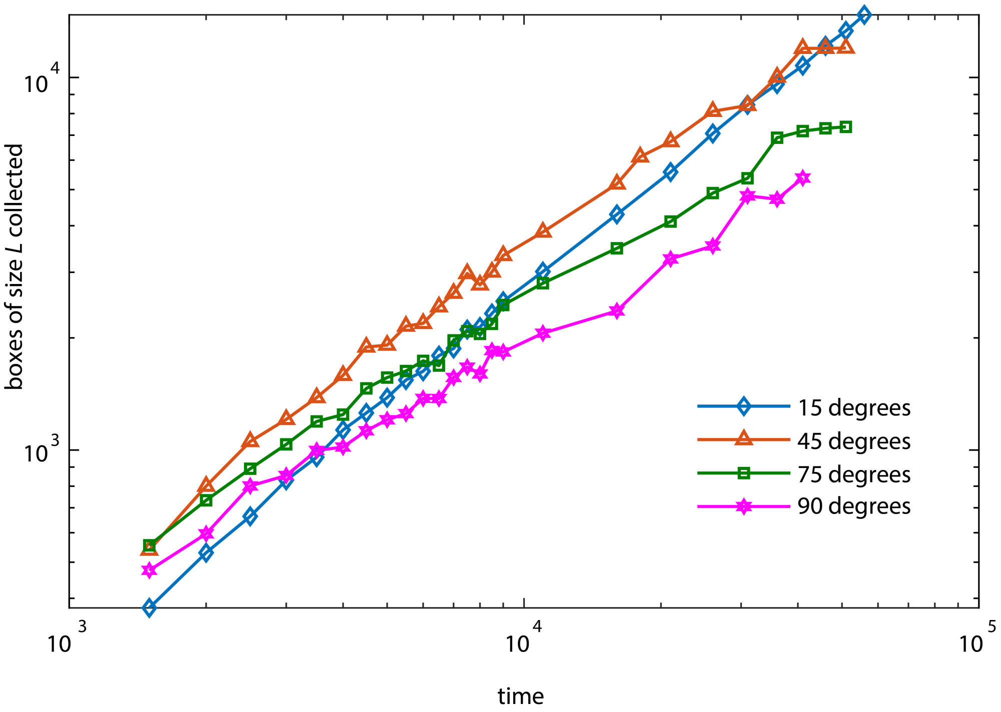

**Figure S17: Optimal branching angle changes with search duration.** Otherwise identical networks with four different branching angles were simulated across time. Box counting was used to quantify the number of boxes collected of size $\bar{L}$. Networks collected more boxes with time but sublinearly with different branching angles performing better at different times. The data presented are the average of 10 simulations for each time point. Network parameters: $S = 300$, $k_b = 0.01$, $k_g = 0$, $k_r = 1$

Here we study how the branching angle, $\alpha$, affects the area searched over time. We ran simulations for $\alpha = 15°, 45°, 75°$, and $90°$ and counted boxes of size $\bar{L}$ for the entire history of each simulation (see supplementary section 8.3). The network with optimal $\alpha$ for search at this resolution is the one that collects the most boxes.

We find that over short times the network with low $\alpha$ of $15°$ performs worse than networks with larger $\alpha$ at this search resolution. However, as time progresses the optimal $\alpha$ for this search task decreases until the network with $\alpha = 15°$ performs best (figure S17). These networks have the same rates and same structure, except for their branching angles, hence the length explored by each network over time should be the same (eq. (S99)). The branching angle has two primary effects on the network. 1) As discussed in section 5.2.1, lower $\alpha$ increases the persistence length. Larger persistence length leads to a slower self-crossing of the network path, as shown in main paper figure 3. Self-crossing has the effect of searching redundant boxes which do not newly contribute to the count. Thus networks slower to self-cross continue searching fresh area for a longer duration. 2) $\alpha$ also impacts the characteristic network width and thoroughness of search as discussed in section 6.1.1. Too small an $\alpha$ leads to a network that does not thoroughly search the area away from the path. Networks with intermediate $\alpha$ with $w \sim L_p$ leave fewer areas along the path unexplored as they advance. The optimal balance between these two competing impacts of alpha on search shifts over time, with large persistence lengths (low $\alpha$) favored over longer times as self-crossing becomes more and more common, and $w \sim L_p$ favored at early times before self-crossing becomes dominant.

##### 6.1.6 Optimizing search

Finally, we point out that whether any of these search behaviors (and corresponding parameter choices) are optimal ultimately depends on the search goal, for example, performing a coarse search over a large area, or performing a thorough local search with significant self-crossing of the network. As shown above, the optimal parameter choices also depend on the network size and duration of the search. We note that various aspects of search, such as self-crossing and the ability to reach a given point in space also depend on the dimensionality of the space the network is embedded in.

#### 6.2 Active search

Having gained an understanding of the search implications for the unbiased stochastic movements of networks with different parameters, we next focus on how bias can be introduced to cause directed travel in landscapes of varying environmental quality, $E$. Making some or all of the rate constants for individual leaves depend on $E(x)$ at the specific position $x$ where each leaf resides can result in biased motion. Here we discuss some of the more general features of biased travel and their implications for active search.

##### 6.2.1 Network strategies for biased motion

The number of free leaves grows according to eq. (S2), $\frac{dN_F}{dt} = k_b N_F - k_s N_F$, thus the difference, $\delta = k_b - k_s$ determines the rate of change of the number of free leaves, increasing exponentially with $\delta > 0$ or decreasing exponentially with $\delta < 0$. In the unbiased, steady state model, $k_b = k_s$, such that the network maintains a constant number of free leaves.

Modifying $\delta$ provides one way to bias the network motion. This can be accomplished by changing either $k_b$, $k_s$, or both. If $\delta$ is different at different locations within the network, these locations will experience different growth rates in the number of free leaves. It is especially effective to make $\delta$ positive in some areas (causing exponential increase of those areas) and negative in other areas (causing exponential decrease of those areas), resulting in net movement in the direction of larger $\delta$. If rates are changed in such a way to make $\delta$ a function of the environment the network can be made to move toward (or away from) particular environmental conditions, for example ascending (or descending) an environmental gradient.

Furthermore, since the unmodified model is at the critical point between expanding and shrinking, only small changes to $\delta$ are needed to introduce substantial bias, since the branching nature of the model automatically amplifies the number of free leaves exponentially according to $\delta$. Hence, maintaining the model at the critical point between exponential growth and decay allows the network to have sensitive response based on small changes to $k_b$ and $k_s$; changes in rate that are only linear with environment can bring about exponential responses of the network. Changes to other rates such as retraction or growth, $k_r$ or $k_g$, do not directly amplify exponentially in this manner, thus larger changes to these rates are needed to bias the network's travel.

We implemented two different schemes to effect network response to an overlaid environment. 1) In the first scheme, $\delta$ is modulated for each free leaf according to the comparison of its environment to the mean environment across all free leaves. Averaging over all leaves has the effect that the network self-tunes to stay consistent in response across arbitrary absolute environmental conditions, and arbitrary slopes of environmental condition. In other words, comparison to the mean environment across leaves keeps the network at the critical point and maintains sensitive response across different environments. Other types of average, such as the median, may be more favorable, depending on environment. 2) In the second scheme, each free leaf responded locally to its environmental conditions without regard to conditions of other leaves by modifying its restructuring rates. This scheme doesn't allow equal response across arbitrary absolute environmental conditions and environmental gradients, however, this scheme also does not require that leaves share and integrate information about the environment with other leaves (which may not always be possible). Both schemes bias the direction of network travel. More detail on the specific implementations for these two schemes can be found in section 8 and examples in supplementary movies 5-9.

### 7 Alternative traveling network models

#### 7.1 One-state 'uncommitted retraction' traveling network model

a

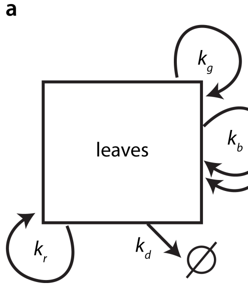

b

| | $k_b$ | $k_g$ | $k_r$ | $k_d$ |
|---|---|---|---|---|
| $S$ | +2 | +1 | -1 | 0 |
| $N_1$ | +1 | 0 | 0 | -1 |
| $N_2$ | 0 | 0 | 0/-1 | +1 |
| $N_3$ | +1 | 0 | 0 | -1 |

**Figure S18: State diagram and table of restructuring impacts for single state network model.**

We also considered a simplified version of the original model with only one leaf type and no switching events. This has the effect that leaves can go back and forth between growing and retracting. Figure S18 illustrates the corresponding rates and the conversion tables for $\Delta L = 1$.

Equations for this one-state model in steady state (for $\Delta L = 1$):

$$\begin{aligned}
\frac{dS}{dt} &= 0 = N_1(2k_b + k_g - k_r) \implies 2k_b + k_g = k_r \\
\frac{dN_1}{dt} &= 0 = N_1(k_b - k_d) \implies k_b = k_d \\
\frac{dN_3}{dt} &= 0 = N_1(k_b - k_d) \implies \text{redundant relation} \\
\frac{dN_2}{dt} &= 0 = N_1 k_d - N_1 k_x \implies k_x = k_d = k_b.
\end{aligned}$$

Our simulations show that regardless of size and parameters, networks under these constraints develop into a single nearly unbranched chain of degree-two nodes (supplementary movie 10). The motion of this one-state model also becomes diffusive but has much lower diffusivity than the model with irreversible retraction for the same growth and branching rates. Diffusivity also decreases in this one-state model for larger network sizes (figure S19), whereas in the original 'committed-retraction' model diffusivity is independent of network size. In other words, including the simple rule – 'finish retraction once started' – results in more effective motion, invariant long-term diffusion with size, and a greater diversity of possible structures. This may help explain why, from our observations, 'leaves' of moving *Physarum* typically do not begin extending again once they have started retracting (supplementary movies 1 and 9), and why decisive actions in a businesses (stick with your decision to retract) might be a good strategy. This aspect of the two-state model is also somewhat reminiscent of actin treadmilling where one end of a filament retracts continuously while the opposite end branches and grows (*12*).

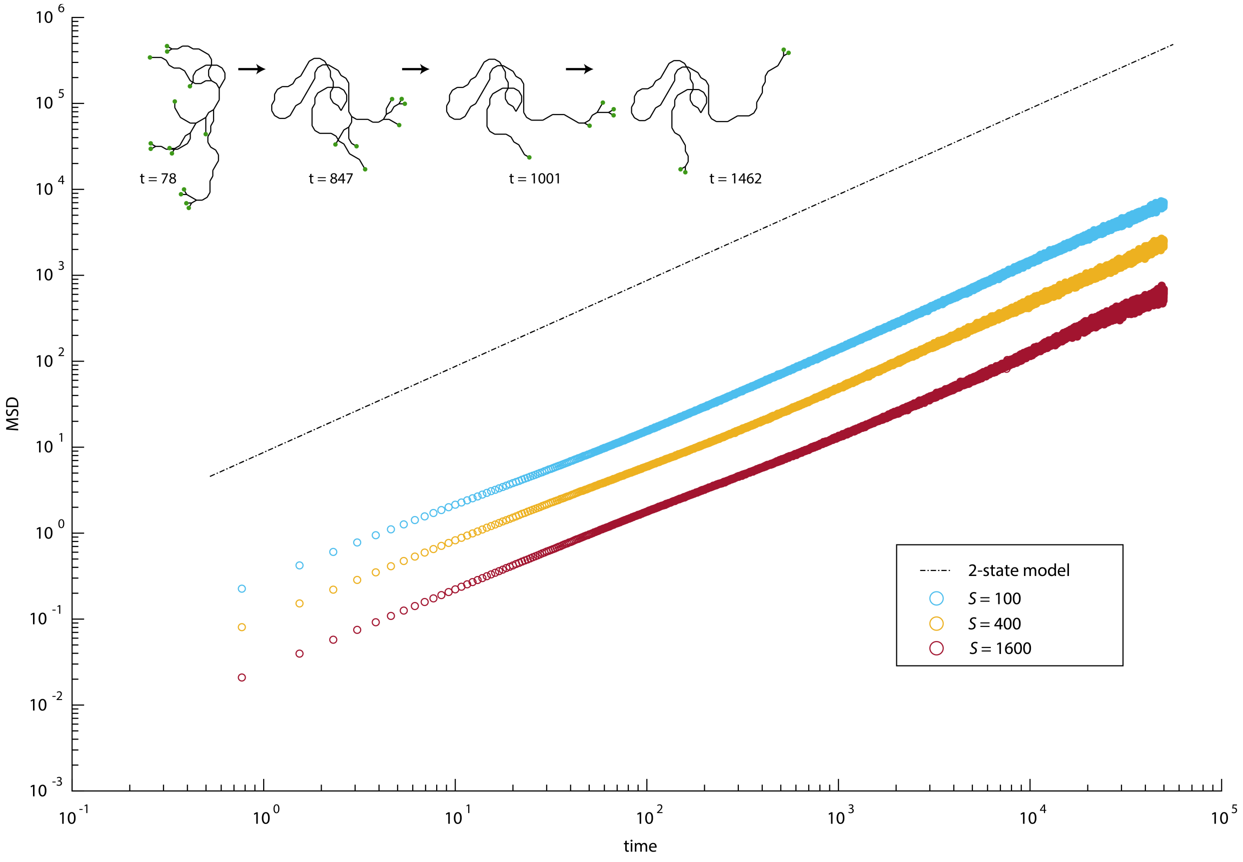

**Figure S19: Mean squared displacement vs time for three different network sizes with reversible retraction.** Networks with reversible retraction diffuse much slower than those with irreversible retraction. Larger networks with reversible retraction diffuse slower than smaller networks with reversible retraction. Network parameters $k_b = 0.2$, $k_g = 0.1$, $k_r = 0.5$, $\alpha = 60$ over $\Delta t = 10,000,000$ shown here. The dashed black line is the analytical diffusive prediction from the model with irreversible retraction for the same $k_b$ and $k_g$, showing much faster diffusive motion. Inset images show a representative progression to a single chain of nearly-unbranched degree-two nodes for a network of size $S = 100$, same parameters as above, leaves indicated in green.

#### 7.2 Other traveling network model possibilities

Many other related traveling network models remain to be explored, both for understanding other theoretical aspects as well as better capturing specific real-world systems. We briefly mention several we consider particularly interesting:

(1) Instead of distinct free and retracting leaf states, all leaves could be allowed to retract, grow, or branch. This leads to marked differences (see previous section 7.1) which highlight the utility of the two state model analyzed in the paper.

(2) Nodes of higher degree could be allowed, by, for example, not adding any edge length during branching.

(3) Dynamic rules could be implemented to allow for the formation and breaking of loops, i.e., for cyclic networks. This model would allow networks that stay more robustly connected if subjected to random edge or node annihilation. Under certain assumptions, connective shortcuts (analogous to those in small-world networks (*13*)) might allow more efficient redistribution of network resources and faster network relocation. Under steady state, this model would require another 'topological' condition to be met, and the network would no longer always follow a single unique path. This might better capture large *Physarum* organisms which can form and break loops.

(4) The individual edges might carry different weights. This could be used to capture scenarios when resource cost of connectivity is not always strictly proportional to the edge length. For example, in *Physarum* different connections can have different widths to optimize transport (*14*).

(5) The network size $S$ could vary over time, e.g., for physarum a steady fractional decrease in $S$ would mimic energy expenditure, while the collection of resources (food) would lead to an increase. This could be easily coupled to time varying environments that either change on their own or as a result of network activity as in main paper figure 4a and 4b. Investigating dynamic changes to the network size may expose additional insights not revealed by our primarily steady state analysis of the model.

(6) Internal and edges nodes could be manipulated. This might allow networks to more efficiently structure themselves and allow more flexibility in how structure and motion are coupled. Specific changes, such as allowing branching off of established edges, would better capture the actin network, where branching does not happen exclusively at the leaves.

(7) Considering the embedding of networks in higher dimensional spaces. Many aspects of these networks are independent of dimension, but questions of search and network shape may become interesting to think about, and the abstract spaces inhabited by organizations are certainly many-dimensional.

(8) Memory and self-avoidance are doubtless used by real systems to allow more efficient movement and search. For example, *Physarum* deposits a trail of extracellular slime proposed to provide spatial memory (*15*) and enhance foraging (*16*).

### 8 Numerical implementation of the model

We implemented the model presented in the main text with a stochastic simulation.

#### 8.1 Base simulation

We created a base simulation upon which we added additional features. Each network node was stored with an identifier, its connected partners, location coordinates, and state (free or retracting for leaves). According to the model described in the main text and supplementary section 3, leaves were manipulated to generate the network dynamics according to branching, growth, switching, and retraction according to their associated rates.

Time was discretized in the simulation by $dt$ to give probabilities for each action over dt of $P_b = k_b \cdot dt$, $P_g = k_g \cdot dt$, $P_s = k_s \cdot dt$, and $P_r = k_r \cdot dt$. The time increment $dt$ was chosen such that the sum of probabilities for all actions of any leaf was never greater than 1. As described in the main text and supplementary section 3: Growth displaces the associated node a distance $\Delta L$ away from its connected partner. Branching creates two new leaves, each distance $\Delta L$ from the original node at angle $\alpha$ from each other. The branching angle, $\alpha$, was included as a tunable parameter. Retraction displaces the associated node a distance $\Delta L$ toward its connected partner. If the partner node of a retracting node is reached, the leaf is eliminated and the retracting state is passed to the partner node if the retracting node becomes a leaf.

##### 8.1.1 Maintaining steady state

As derived in section 4.1.1, setting $k_s = k_b$ is necessary to maintain a steady state number of free leaves and, by eqs. (S35) and (S36), network size, but is itself not a sufficient condition for maintaining network size in a stochastic implementation of the model. Without an additional constraint the mean size across many networks would remain the same, but individual network sizes would change according to a drift process. For steady state simulations, in addition to setting $k_s = k_b$, we also included a Hill function that modified the switching probability to $P_{smod}$ to act as a constraint pushing the size toward a set size, $S_{eq}$.

$$P_{smod} = \frac{2P_s}{1 + (\frac{S_{eq}}{S})^n}, \tag{S100}$$

where $S$ is the network size and $n$ is the Hill coefficient. This function causes the switching rate to increase if the network becomes larger than the set size (which has the correcting effect of decreasing the size) and the switching rate to decrease if the network size is less than the set size (which has the correcting effect of increasing the size).

We included the Hill coefficient as an optionally tunable parameter in the simulations but held it constant at $n = 10$ for all results presented here. Changes to $n$ did not seem to change the long time average structure and motion of the simulation. The remaining rates were held constant across dynamic size fluctuations, and, over long time scales, the average switching rate will still be equal to the branching rate.

Despite the modified switching probabilities from the Hill function, it could still be possible for all free leaves to switch to the retracting state. This would result in a situation with no free leaves and a collapsed network (since retracting leaves don't switch back to the free state in this model). Networks with few free leaves are more prone to this trivial outcome. To avoid this possibility, we included a rule enforcing that the last two free leaves cannot switch to the retracting state, even if their stochastic switching probabilities would have had them do so. This guaranteed that the networks never collapsed during simulations of arbitrary length.

##### 8.1.2 Advancing through time

The simulation then stepped through time to some desired $\Delta t$ in $dt$ increments to generate the network dynamics. At each time point, each leaf was sequentially manipulated according to its allowed actions with corresponding probabilities and updated accordingly. The size of the network was tracked with each action and $P_{smod}$ was also updated after each action. Edges can optionally be plotted to visualize the model using the stored partner and spatial coordinate information.

The result of this simulation was a network that dynamically restructures itself and travels through space. Example simulation outputs are shown in supplementary movies 2 and 3. The simulation was used to gain an intuitive understanding of this traveling network model, generate visualizations of typical networks, and compare to analytical results.

Note that for simplicity and correspondence to the analytical results, the model was not self-avoiding, and the spatial embedding was performed in two dimensions, but as noted in the text, many structural aspects of the model do not depend on the dimensionality. This simulation was implemented in MATLAB. Additions and modifications to the model are discussed in the relevant sections.

#### 8.2 Simplified model for limit case of $\bar{L} = 1$, $k_r \gg k_b$

We also implemented a simplified version of the model to investigate the limit case of $\bar{L} = 1$, $k_r \gg k_b$. This model was simulated according to the simple rules discussed in section 4.4.7: i) Begin with a binary tree. ii) If the total network size exceeds the threshold size, $S$, then randomly select a leaf and fully retract it past all degree-two nodes until a degree-three node is reached or only two nodes remain. iii) Randomly select a leaf and branch it. iv) Return to ii. As noted in section 4.4.7, retraction events in such a model are scale free such that there is a small, but finite probability of the entire network retracting such that no edges remain. This corner case is addressed in step ii by constraining the model to stop such retraction events while one edge still remains (i.e. before complete network collapse). This imposes a limit on the size of the maximum relaxation event, analogous to the limit set by the system size in classic SOC sand pile models (figure S9) (*6*). Note that this model undergoes large transient reductions in size after large retraction events.

#### 8.3 Fractal dimension and box counting:

To determine various aspects of network shape and search, we used the box counting method from fractal or Hausdorff dimension to quantify the area covered (*11*). The full network history (all nodes over all simulated time) was saved and converted into an image file with edges rendered as lines. Care was taken while saving these images to assure the absolute image scale and line width to pixel ratio remained constant to allow meaningful comparison between results across different simulation input parameters (since different $k_b$, $k_g$, network size and runtimes result in networks whose trajectories have vastly different absolute sizes).

Boxes of a given size were then generated and tiled across the image. The total number of boxes containing a network pixel were counted, and the tiling and counting was repeated for numerous box sizes to generate the box counting results (*11*).

Networks were initialized as a single short edge at $t = 0$ and expanded over time to equilibrium sizes. To avoid confounding effects we waited until after the transient initial network growth to begin recording the network history.

#### 8.4 Environmental response

Many possible schemes allow this traveling network model to respond to it's environment. We implemented two, as shown in supplementary movies 5-9 and main paper figure 4 and as discussed below.

##### 8.4.1 Environmental response: Scheme 1

We implemented one scheme by modifying the switching probability for each free leaf according to a comparison of that leaf's local environment, $E_i$, to the mean environment across all free leaves, $\bar{E}$.

We created an environmental resource field upon which the networks exist and travel. Distribution and abundance of resources in this field can be set to any arbitrary function. We used both Gaussian peaks and linear gradients in the main text.

The mean environmental condition, across all free leaves $N_{f1}$ to $N_{fn}$ was computed by averaging the value of the environmental field for all free leaves $\bar{E} = \frac{1}{n}\sum_{i=1}^{n} E_i$, then setting the final switching probability, $P_{sEmod}$, equal to 0 for all leaves with local environmental values greater than $\bar{E}$, and $2P_{smod}$ for all leaves with local environmental values, less than or equal to $\bar{E}$. For each free leaf,

$$P_{sEmod} = \begin{cases} 0 & \text{for} \quad E_i > \bar{E} \\ 2P_{smod} & \text{for} \quad E_i \leq \bar{E} \end{cases} \tag{S101}$$

In this response scheme the mean environmental condition was computed once per time point. Optionally, noise can be added to the environmental reading for each leaf, for example, Gaussian noise for each leaf, at each time point, as in main paper figure 4c. This type of scheme allowed for response to arbitrary values of the environmental field magnitude and slope (excluding noise).

We also implemented a two-way interaction with the environment where the network responded to and also modified its environment. Response of the network to the environment was simulated as above. The response of the environment to the network was implemented for each Gaussian hill in the environment according to the presence of free leaves. Each free leaf subtracted from the peak according to that peak's contribution to the leaf's environmental value multiplied by a 'feeding rate'. In this way, if free leaves were located on a peak, the height of that peak decreased as seen in supplementary movie 6 and main paper figure 4b.

##### 8.4.2 Environmental response: Scheme 2, local response

Scheme 1 required sharing information about the environment between leaves, which may not be possible or practical for all distributed systems. We also created a simulation where the network acted on environmental information that was local to each leaf (scheme 2).

We implemented this scheme by modifying the branching, growth, and switching probabilities according to:

$$P_{bEmod} = P_b + P_b \cdot E_{local} \tag{S102}$$

$$P_{gEmod} = P_g - P_g \cdot E_{local} \tag{S103}$$

$$P_{sEmod} = P_{smod} - P_{smod} \cdot E_{local} \tag{S104}$$

where $E_{local}$ was the environment local to each free leaf. Here we implemented the environmental resource field slightly differently than above. Instead of continuously defined functions, we created a pixelated resource field where each pixel had a value between 0 and 1 and $E_{local}$ for each leaf was the pixel overlapping the spatial coordinates of that leaf.

We included response of the environment to the network by having each free leaf 'consume' its corresponding environmental pixel according to a feeding rate, $R_f$. In this scheme we also implemented dynamic changes to the network size, $S_{eq}$. As the network consumed resources, we increased $S_{eq}$ according to the feeding rate multiplied by a size gain rate, $S_g$, as, $\frac{dS_{eq}}{dt} = S_{eq} + R_f \cdot S_g$. The network also lost equilibrium size according to a shrinkage rate, $S_r$, as a fraction of the total equilibrium size, $\frac{dS_{eq}}{dt} = S_{eq} - S_{eq} \cdot S_r$, which could represent, for example, the basal resource cost associated with existing and moving.

The results of this implementation are shown in supplementary movie 7, which shows an example network with initial $S_{eq} = 75$, $k_b = 0.2$, $k_g = 0.45$, responding to and modifying a randomly seeded environment with a feeding rate of $R_f = 0.25$, size gain rate $S_g = 1$, and shrinkage rate $S_r = 0.005$. This second implementation demonstrates that there are multiple ways to generate responses in these networks, and the environmental response and modification scheme should be chosen to best capture particular systems and properties of interest.

### 9 Cultivation and analysis of *Physarum*

*Physarum polycephalum* was cultivated as in Hossian *et al* (*17*). Briefly, *P. polycephalum* (Carolina Biological Supply) was maintained on oat flakes and inoculated onto 2% non-nutrient agar (BD Bacto Agar). An HP Scanjet G3110 flatbed scanner was used to record the behavior, taking images every 10 min. In supplementary movie 9 food was deposited as droplets of oatmeal water with a custom robotic setup and indicated by a digital overlay, as described in Hossian *et al* (*17*).

Image analysis for displaying the 'age' of different parts of the organism in supplementary movie 1 was performed by subtracting the background from images, then extracting the pixels representing the organism from these images using the 'ilastik' pixel classifier (*18*) trained on 19 labeled physarum images. Pixel age as in supplementary movie 1 was defined as the number of frames that a particular pixel had been occupied. The organism path was extracted manually.

### 10 Supplementary movie descriptions

**Supplementary movie 1:** The *Physarum polycephalum* network travels through space during foraging behavior. The movie shows a single organism traveling on the flat surface of 2% agar without food or nutrients for 24 hours. The second portion of the movie replays the same sequence where the organism shape has been extracted from the background and each pixel colored according to the length of time the organism occupied it. The path that the organism took through its environment is displayed in gray. Each frame is 10 minutes apart, organism is $\sim$1 cm, scale bar given in main paper figure 1. See supplementary methods for information on the experimental details.

**Supplementary movie 2:** Results of simulating the traveling network model presented in the main text with size 60, $K_B = 0.2$, and $K_G = 0.1$. Free leaves are colored cyan, and retracting leaves are red. Degree -two and -three nodes and edges are black. See supplementary methods section 8 for details of the numerical implementation. Time and length are arbitrary units.

**Supplementary movie 3:** Results of simulating networks of size 70 with various $K_B$ and $K_G$ values. Free leaves are colored cyan, and retracting leaves are red. Degree -two and -three nodes and edges are black. Here the frame pans to keep each network centered as they travel and, time and length are arbitrary units.

**Supplementary movie 4:** Illustrates head, path, and tail concepts. First a traveling network with size 20, $K_B = 0.2$, and $K_G = 0.1$ is simulated with node color indicating the node age. The clip is then rewound and played again with the tail (oldest node, red), head (leading node on path, red), and path (black line) indicated. Time and length are arbitrary units.

**Supplementary movie 5:** Shows a traveling network responding to a Gaussian hill of resources in its environment. The response is implemented with the scheme discussed in supplementary methods section 8.4.1 and, time and length are arbitrary units.

**Supplementary movie 6:** Shows the traveling network responding to two Gaussian hills of resources in its environment and the environment, in turn, responding to the network. The responses are implemented with the scheme discussed in supplementary methods section 8.4.1 and, time and length are arbitrary units.

**Supplementary movie 7:** Shows a traveling network responding to a random resource field. In this response scheme leaves do not share environmental information, but instead modify rates according to local environmental information according to the scheme discussed in supplementary methods section 8.4.2. Time and length are arbitrary units.

**Supplementary movie 8:** Shows one or three traveling networks responding to one or three Gaussian resource hills. The response is implemented with the scheme discussed in supplementary methods section 8.4.1. The sum of the network sizes in each clip is 150. Multiple smaller networks are able to capture multiple opportunities in more rugged environments whereas a single large network captures only a single opportunity. Time and length are arbitrary units.

**Supplementary movie 9:** Part 1: Shows a *Physarum polycephalum* organism following a trail of food (robotically deposited oatmeal solution) on 2% agar in a 150 mm diameter Petri dish. Red circles are digitally overlaid to show position and timing of food deposition. Each frame is 10 minutes apart, total duration $\sim$2 days.

Part 2: Shows the traveling network model responding to and consuming a trail of resources. The response is implemented with the scheme discussed in supplementary methods section 8.4.1 for a network of size 300, $K_B = 0.05$, and $K_G = 0$, and time and length are arbitrary units.

**Supplementary movie 10:** Shows the single-state traveling network model. Traveling networks under these assumptions degenerate into a single nearly-unbranched chain of degree-two nodes with reduced diffusion compared to the two-state model. Network parameters $k_b = 0.2$, $k_g = 0.1$, $k_r = 0.5$, $\alpha = 60$ over $\Delta t = 1500$ shown here.
